# Taxonomy of neural oscillation events in primate auditory cortex

**DOI:** 10.1101/2020.04.16.045021

**Authors:** Samuel A Neymotin, Idan Tal, Annamaria Barczak, Monica N. O’Connell, Tammy McGinnis, Noah Markowitz, Elizabeth Espinal, Erica Griffith, Haroon Anwar, Salvador Dura-Bernal, Charles E Schroeder, William W Lytton, Stephanie R Jones, Stephan Bickel, Peter Lakatos

## Abstract

Electrophysiological oscillations in the brain have been shown to occur as multi-cycle events, with onset and offset dependent on behavioral and cognitive state. To provide a baseline for state-related and task-related events, we quantified oscillation features in resting-state recordings. We used two invasively-recorded electrophysiology datasets: one from human, and one from non-human primate auditory system. After removing incidentally occuring event related potentials, we used a wavelet transform based method to quantify oscillation features. We identified about 2 million oscillation events, classified within traditional frequency bands: delta, theta, alpha, beta, low gamma, gamma, high gamma. Oscillation events of 1-44 cycles were present in at least one frequency band in 90% of the time in human and non-human primate recordings. Individual oscillation events were characterized by non-constant frequency and amplitude. This result naturally contrasts with prior studies which assumed such constancy, but is consistent with evidence from event-associated oscillations. We measured oscillation event duration, frequency span, and waveform shape. Oscillations tended to exhibit multiple cycles per event, verifiable by comparing filtered to unfiltered waveforms. In addition to the clear *intra*-event rhythmicity, there was also evidence of *inter*-event rhythmicity within bands, demonstrated by finding that coefficient of variation of interval distributions and Fano Factor measures differed significantly from a Poisson distribution assumption. Overall, our study demonstrates that rhythmic, multi-cycle oscillation events dominate auditory cortical dynamics.

## Introduction

Intrinsic cortical oscillations consist of both rhythmic and brief, pulse-like neuronal activity patterns, which co-occur in electrophysiological recordings [1–3]. These patterns manifest differently across multiple frequency bands and different brain regions during various task-dependent brain states [4]. These nearly-continuous neural oscillations can be viewed as providing a background context on top of which behaviorally and cognitively relevant information (content) is transmitted [5].

Two related theories address the role of oscillations. *Entrainment theory* posits that attentional selection involves phase reset in response to relevant sensory or internal inputs; phase alignment provides optimal response to relevant time-varying signals [1,5–8]. Similarly, *communication through coherence* posits that synchronization of oscillation phase across different brain circuits optimizes the ability of the networks to transmit and receive information [9]. The presence of nearly continuous oscillations at particular frequencies (*e*.*g*., gamma) is important for these mechanisms to work, since this provides the context for *sender* and *receiver* networks to periodically communicate [10–12]. In other words, oscillations can be used to form dynamically changing functional networks based on their matched or mismatched timing and phases. Intrinsic oscillations should be distinguished from stereotypical event related potential (ERP) waveforms, which may also be prolonged enough to produce a single cycle of oscillation.

Delta (0.5-3 Hz), theta (4-8 Hz), alpha (9-14 Hz), beta (15-30 Hz), gamma (30-100 Hz), and higher frequency oscillations have been shown to occur as multi-cycle events, with onset and offset dependent on behavioral and cognitive state [5,13]. A prominent example is the primate alpha rhythm (9-14 Hz) observed in the primary visual cortex [14]. Rhythms in the same frequency range have also been observed in primary auditory, somatosensory, and motor cortices, as well as in subcortical structures [15–19].

The presence of high spectral power in a neural signal does not necessarily indicate an intrinsic oscillation. This is true particularly for one- or two-cycle events which could arise stochastically [2,3,20]. For example, a high-amplitude, single-cycle waveform with 50 ms duration could be mistaken for a 20 Hz oscillation [2,21]. If underlying neural generators are stochastic, high spectral power may simply reflect intrinsic temporal domain features from synaptic time constants or other sources [2,20]. Recent studies that support this interpretation demonstrated recurring brief events in high-frequency gamma [22,23], and in beta [21,24–26] ranges. Therefore, care must be taken to distinguish non-oscillation waveforms due to time-domain features from clear multi-cycle oscillations. The relevance of this dichotomy for particular frequency bands, particular behavioral conditions, and particular brain regions remains to be determined.

Our aim was to examine spontaneous neuronal activity, recorded over long time scales (minutes) in awake, resting state conditions in humans and non-human primates (NHPs), in order to better understand basic features and temporal properties of physiologically-relevant oscillations in auditory cortex. We used two invasively-recorded electrophysiology datasets: 1) laminar local field potential (LFP) recordings from NHP primary auditory cortex (A1), and 2) intracranial electroencephalogram (iEEG) recordings from human superior temporal gyrus (STG). We extracted moderate to high-power oscillation events from wavelet transform spectrograms, and determined their basic properties. These included temporal and frequency span, peak frequency, peak power, number of cycles, number of local peaks in filtered waveforms, correlation between filtered and raw signal (*Filter-match*), and rhythmicity across individual events.

Our analysis revealed clear evidence of multi-cycle oscillation events in all frequency ranges (average: 3-4 cycles per event, range: 0.3-44 cycles per event), which we verified by inspection of unfiltered waveforms. Event occurrence in each frequency band also demonstrated rhythmicity, quantified through an analysis of interevent intervals. Overall, our data and analyses demonstrate several characteristics of oscillation events that may be studied in ongoing or single-trial data. Importantly, we demonstrate that multi-cycle oscillation events are prominent in resting state auditory cortex dynamics, and that these events recur quasi-rhythmically over time.

The paper is organized as follows. First, we present a validation of our method using simulations, followed by specific examples of oscillation events in the time and frequency domains. Then, we summarize several characteristics of oscillation events for each frequency band and their intra-event variability. Finally, we look at shifts of events across different bands and cross-frequency interactions.

## Results

### Validation of *OEvent* package for oscillation event detection

We developed the Oscillation Event (*OEvent)* software package for the detection and analysis of electrophysiological oscillation event features. We validated *OEvent* by measuring its accuracy in detecting the number of cycles and the peak frequency in a simulated dataset of sinusoidal signals in several frequency bands and compared the results with a different oscillation detection method which is based on a power and duration threshold termed Better Oscillation Detection (BOSC) [27,28]. Oscillation length was varied from 1-15 cycles with a sufficient interval between event initiation to allow for event detection. For each band, oscillation event signals were superimposed on an NHP auditory cortex supragranular current source density (CSD) signal to provide realistic physiological background signal statistics (**Fig. 1**).

**Fig. 1.**
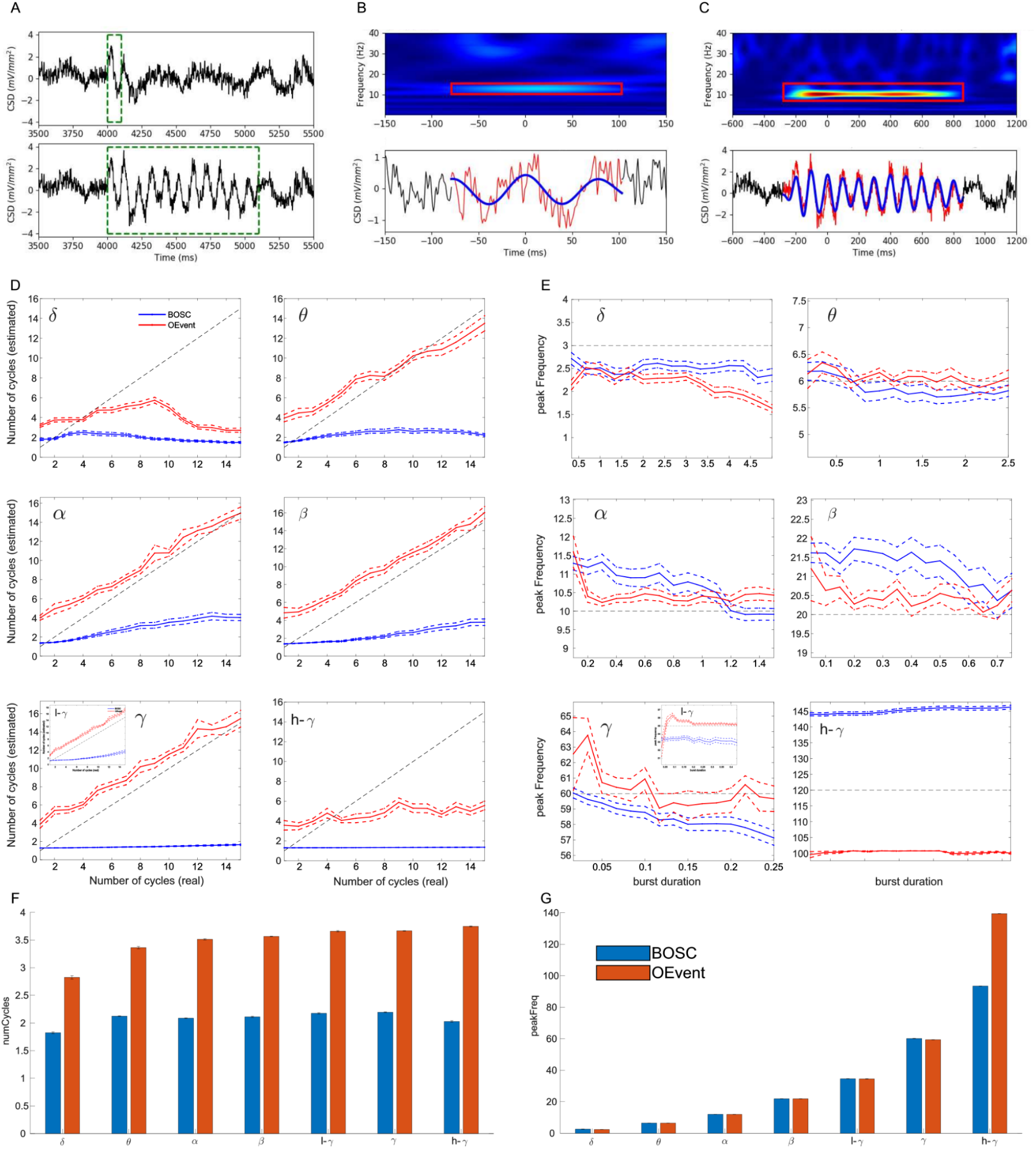
Validation of event detection algorithm. **(A)** Example **of** 1-, 11-cycle simulated 10 Hz alpha signals (green bounding boxes) added to supragranular CSD; amplitude 1.5 mV/mm2. **(B)** 1 cycle signal from **A** was detected as 2 cycles. **(C)** 11-cycle detected as 11.7 cycles. For **B,C** Top: wavelet transform spectrograms with detected event in boxes; bottom: raw (red), filtered (blue) signal **(D)** Detected number of cycles as a function of the actual burst duration (dotted black line) for BOSC (blue) and OEvent (red). **(E)** Peak frequency detected was generally close to the frequency of the simulated signal (horizontal gray line) using both methods in most frequency bands but varied across different burst durations (SEM: dashed lines; insets in D and E show the results for low-gamma frequency band). **(F, G)** Number of cycles and peak frequency detected in NHP A1 recordings using BOSC (blue) and OEvent (red).

As expected, a single cycle alpha event was difficult to detect correctly (**Fig. 1A,B**). In the example shown in **Fig. 1A (top)**, a single cycle alpha event was added to the CSD signal. The random placement of the single cycle event was such that it was followed by an additional cycle of the same frequency that appeared in the CSD data, something that can readily occur randomly in this oscillation rich environment. Unsurprisingly, OEvent detected 2.3 cycles and over-estimated the peak frequency relative to the added cycle (12.75 Hz). An additional factor for overestimation of both frequency and number of cycles were the sharp discontinuity transients at the beginning and end associated with superimposing signal on background.

OEvent was more accurate in estimating the properties of prolonged oscillation events: an 11-cycle alpha event was calculated as 11.7 cycles, with frequency calculated as 10.25 Hz (**Fig. 1A,C**). Estimation accuracy for all frequency bands are shown in **Fig. 1D** (number of cycles) and **Fig. 1E** (peak frequency) for both BOSC (blue) and OEvent (red). In the delta band, both methods struggled to accurately estimate the number of cycles and the peak frequency, specifically for longer durations. At higher frequencies OEvent performed better at estimating the correct number of cycles while both methods performed approximately the same in estimating the peak frequency of the oscillations.

Accuracy across all cycle lengths had an RMS error between 1.45 and 2.46 for theta - gamma bands across all cycle durations. For the delta and high gamma ranges, RMS had higher values of 5.97 and 4.7, respectively, specifically for higher number of cycles (> 6 cycles; **Fig. 1D**). As in the case of **Fig 1A**, in most frequency bands, the number of cycles was typically overestimated for lower-cycle events and performance improved as the number of cycles in the simulated signal increased (except for delta and high gamma bands). Frequency estimation accuracy also varied with the number of cycles (**Fig. 1E**). At a low number of simulated cycles, the peak frequency was more strongly overestimated, which also contributed to overestimation of number of cycles, based on duration and frequency. As in the case of the number of cycles, estimation of the peak frequency had the largest errors in the delta and high gamma ranges. Similar results for the features displayed in **Fig. 1** were obtained when embedding the alpha signal in a background of pink noise (not shown). In addition, we compared the results of the two methods using NHP A1 recordings from two animals (see **Materials and Methods)**. As in the case of the simulated data, the number of cycles detected by OEvent seemed to be higher in all frequency bands and the peak frequency was approximately similar for both methods with a higher frequency detected by OEvent in the high gamma range (**Fig. 1F,G**).

### Characterization of oscillations

8.13 hours of linear array (23 channels spanning 2.3 millimeters encompassing all cortical layers) NHP A1 recordings from 4 animals were converted to current-source density (CSD) time-series to estimate neuronal ensemble transmembrane currents [16]. In this dataset, we detected over 1.9 million putative oscillation events using the OEvent method across all recording channels, after eliminating ERP-like events (see **Materials and Methods**). We also analyzed 37 minutes of human superior temporal gyrus (STG) intracranial electroencephalogram (iEEG) recordings (5 subjects; 43,923 oscillation events). We performed the analyses on the iEEG without calculating a CSD. Re-referencing iEEG signals using a bipolar referencing scheme produced similar results. We focused on characterization of oscillation events, as periods in the wavelet spectrogram with moderate to high power at particular frequencies (defined as above a threshold of 4× median power; see **Materials and Methods**). Events were classified according to traditional frequency bands, with some auditory system specific adjustments: delta (0.5-4 Hz), theta (4-9 Hz), alpha (9-15 Hz), beta (15-29 Hz), low gamma (30-40 Hz), gamma (40-80 Hz), high gamma (81-200 Hz). Event frequency was defined based on the frequency at the point of maximum power during the event.

Our main finding is that oscillations dominated A1 dynamics (**Fig. 2**). Oscillation events in one or more frequency bands were detectable during about 90% of the total recording duration (88.1%, 89.4%, 89.0% for the 3 NHP A1 locations; 89.6% in human STG iEEG). Results were consistent across layers and between NHP and human recordings. Oscillations were typically overlapping or nested.

**Figure 2.**
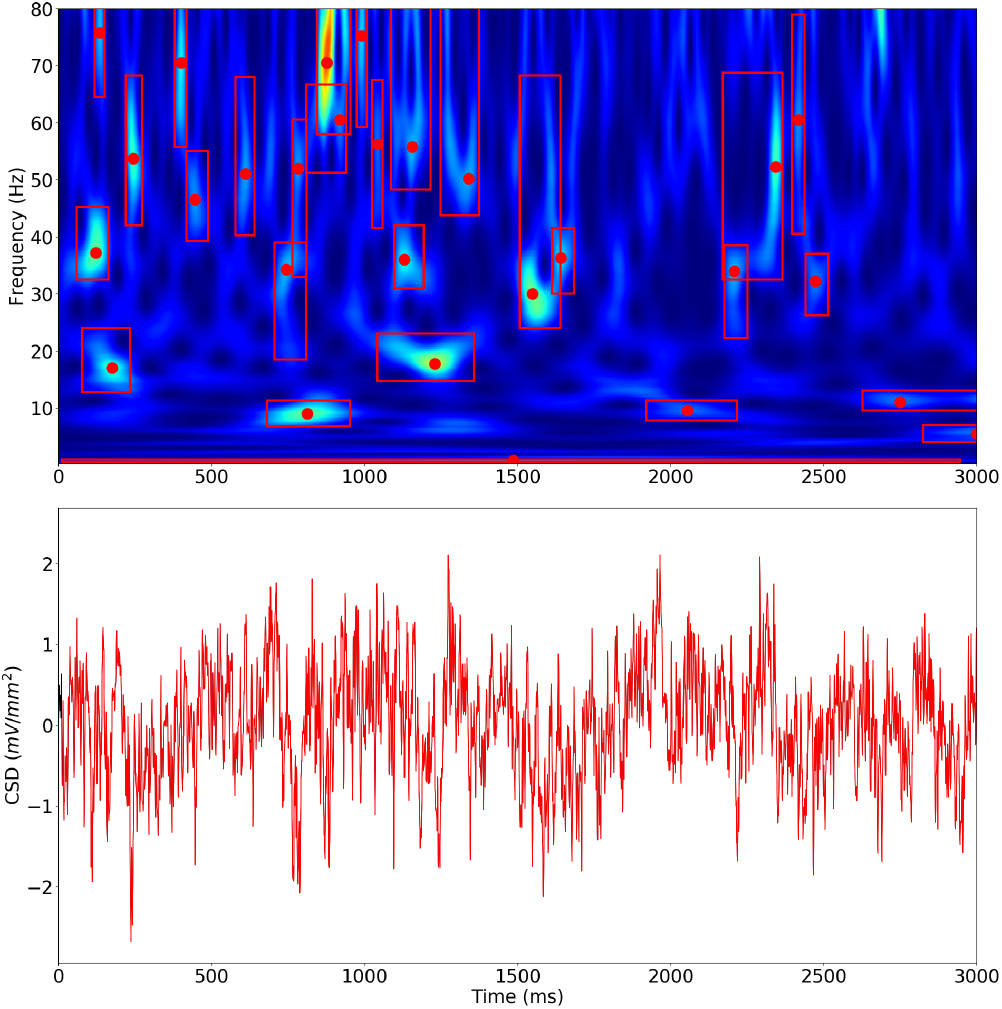
Events occupy the majority of recording time. Example from NHP supragranular A1. Oscillation events (red bounding boxes) occur in one or more frequency bands during this 3 s period. A long delta event is detected from t=0 to near the end of this 3 sec period (red box appears as line across bottom). Red dots show peak frequency; height of box indicates frequency spread. Wavelet transform spectrogram (top) of signal shown at bottom. This example shows events in delta, theta, alpha, beta, and gamma bands.

Figure 3 shows a few examples of oscillation events detected in NHPs data. Oscillation events varied widely in appearance, frequently with crescendo-decrescendo patterns. Many events also showed a pattern of frequency change -- up, down, or U-shaped. For example the alpha event in **Fig. 3** maintained relatively constant power but first increased and then decreased in frequency, as can be seen in the time domain as well as in the spectrum. Despite the relatively broad spread of frequency, this alpha event is clearly continuous, qualifying it as a single oscillation event. The beta event in the same figure showed a different pattern: an abrupt frequency reduction concomitant with power reduction. There was also variation of frequency across events within a frequency band. These observations demonstrate an important aspect of neuronal oscillations that emerges from studying these fluctuations using single-trial (or non-task related) ongoing recordings: oscillation events, even within the same band, might show different characteristics that may shed light on their functional role and the mechanisms generating them.

**Figure 3.**
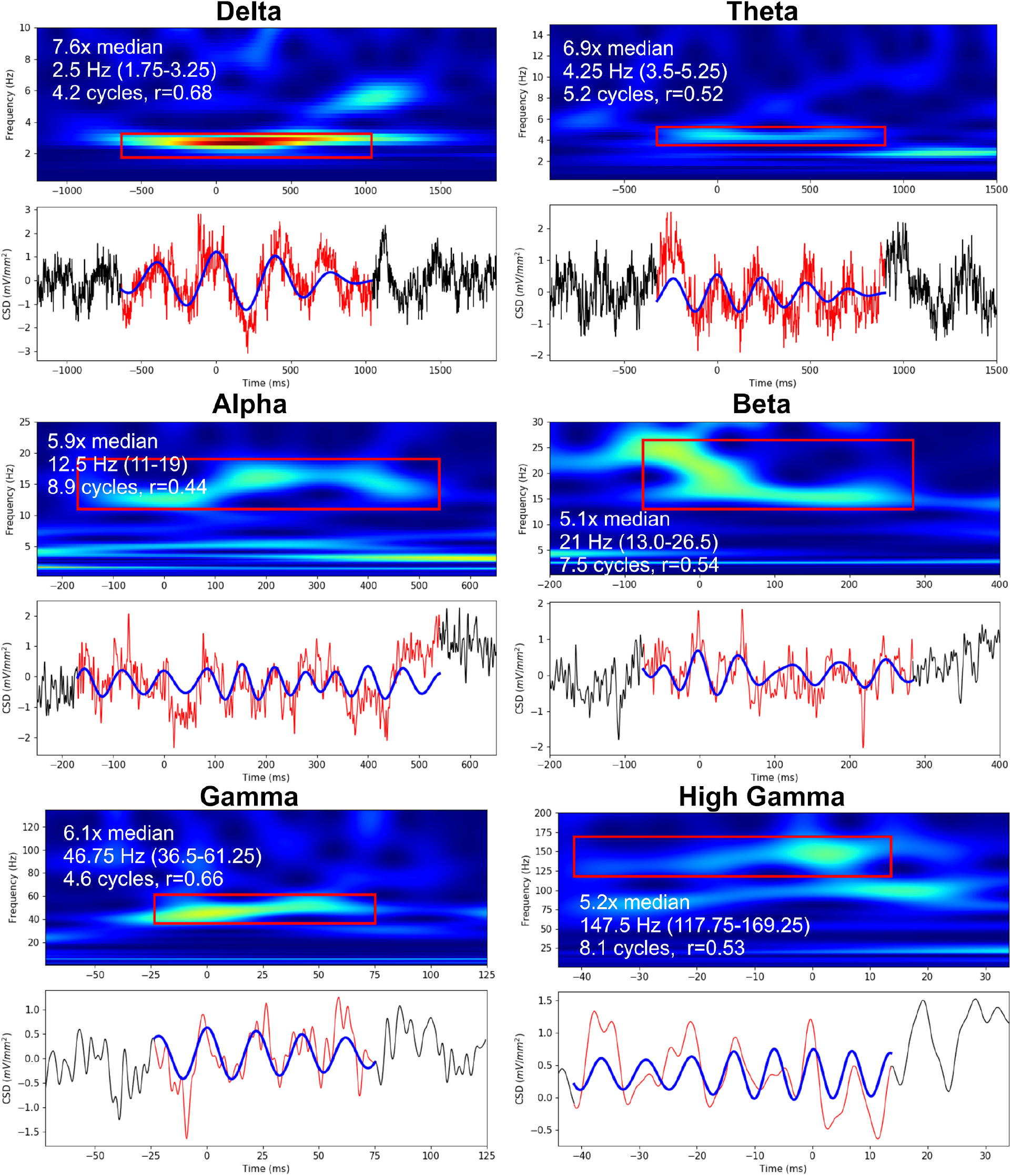
Examples of oscillation events from NHP A1 supragranular layer. Morlet wavelet spectrograms demonstrating individual events (red bounding-box) with raw (red) and filtered (blue) waveforms below (black trace: period outside of detected oscillation). Note that x-y-, and z-(color) scales differ for different bands; power range can be identified from y axis. White text in the spectrograms specifies the events’ power relative to the median, the peak frequency of the event (and frequency range), the number of cycles, and the correlation value between the raw and filtered waveforms (filter-match).

As in prior studies [5,16], delta rhythm dominated in power as well as in “active time” (proportion of the recording time that delta rhythms were present). In both the delta and theta cases, a typical nesting of fast oscillations within the slower oscillation was seen, explaining the relatively low filter-match (correlation between filtered and raw signal; see **Materials and Methods**).

The characteristics of human oscillation events were similar to those of NHP (**Fig. 4**). Oscillation events in the human iEEG were clearly detected across all physiological oscillation bands, with multiple cycles and strong correspondence between raw and filtered waveforms. As with the NHP results, events showed intra-event shifts in frequency and amplitude, for example the alpha event in **Fig. 4** shows a central frequency dip.

**Figure 4.**
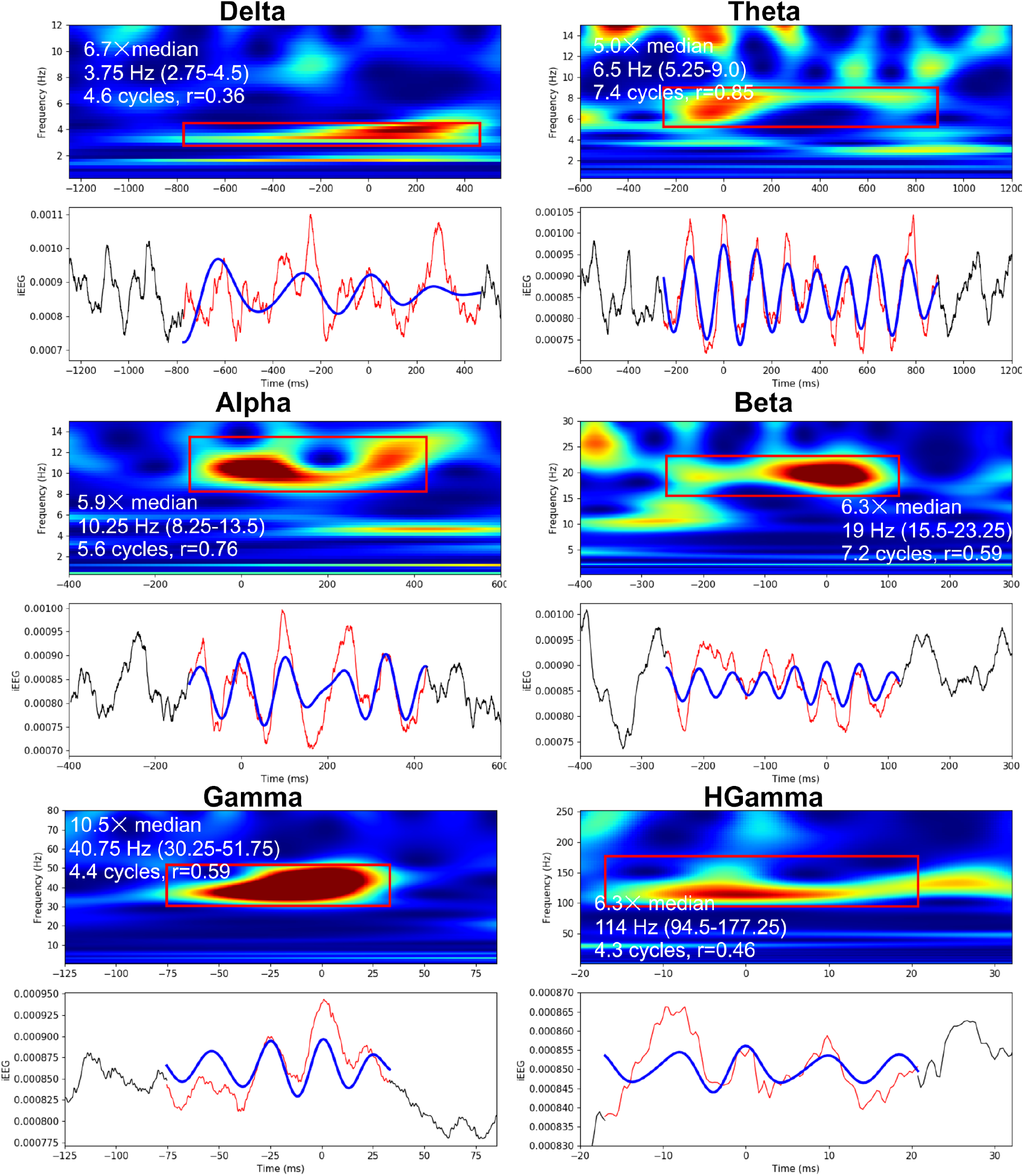
Examples of oscillation events from human STG iEEG. Morlet wavelet spectrograms demonstrating individual events (red bounding-box) with raw (red) and filtered (blue) waveforms below. (x-y-, and z-(color) scales differ for different bands; power range can be identified from y axis; iEEG time-series values are in units of Volts) Time of 0 ms corresponds to the wavelet phase of 0 radians (local maxima) closest to the event’s peak power at threshold detection. White text in the spectrograms specifies the events’ power relative to the median, the peak frequency of the event (and frequency range), the number of cycles, and the correlation value between the raw and filtered waveforms (filter-match).

To better illustrate the time-domain variability in waveform and duration of oscillation events across different bands, **Fig. 5** shows examples of raw and filtered data from NHP A1 neuronal activity sorted by frequency band, with events organized from top-to-bottom by decreasing number of cycles, and left-to-right by decreasing filter-match (Pearson correlation between an event’s filtered and raw signals; see **Materials and Methods**). Oscillations are easier to visually detect in the case of high filter-match. Direct inspection of many such examples allowed us to verify that the algorithm detected “reasonable-looking” oscillations. The oscillations shown here had as many as 32 local peaks (**Fig. 5F**, 2 examples in green bounding box). Combinations of oscillations which co-occurred within a specific event produced differences in waveforms (*e*.*g*, **Fig. 5A**, green bounding box). Importantly, the variability in waveform shapes and oscillation event properties suggests different circuit mechanisms for the production of individual events and might also account for different internal cognitive processes associated with oscillation events within a specific band.

**Figure 5.**
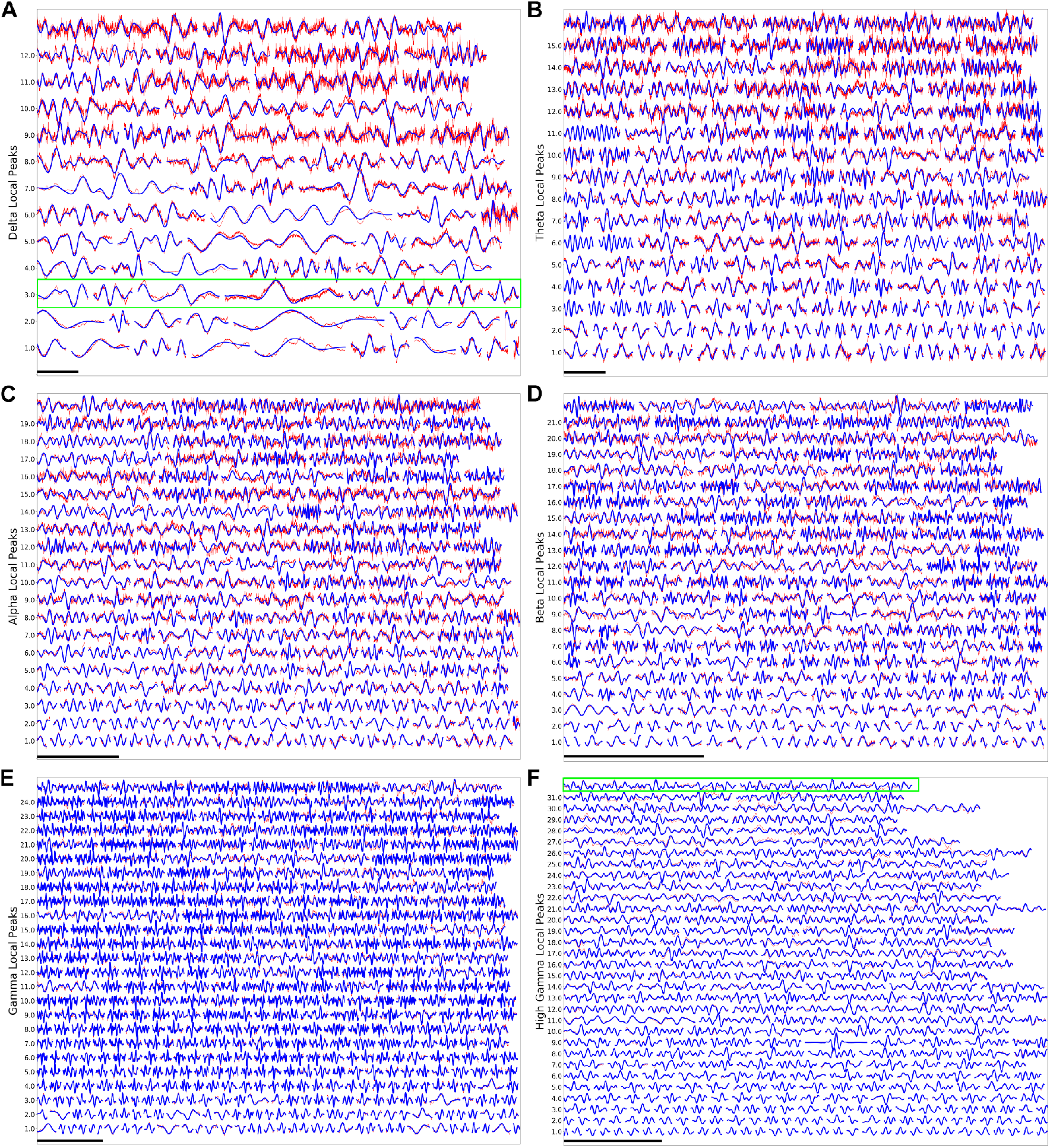
Individual oscillation events from NHP A1. **(A)** Delta, **(B)** Theta, **(C)** Alpha, **(D)** Beta, **(E)** Gamma, **(F)** High Gamma. In each case, decreasing numbers of cycles are shown top to bottom; decreasing filter-match between raw and filtered signal from left to right. Scale bars 1 s except: E: 200 ms; F 100 ms. Examples in green bounding boxes in **(A)** and **(F)** are described in the text.

### Oscillation event features and variability

Next, we quantified the rate of oscillation events at each frequency band and their overall proportion in the recording duration (active time ratio; ATR) for all NHPs and human patients. Lower frequency events will tend to occur more rarely due to their longer duration. Therefore, it was not surprising that higher frequency events occurred most often (**Fig. 6A**). However, longer event durations for delta oscillations produced the longest overall ATR for the delta band (**Fig. 6B**). ATR values ranged from 0.13 to 0.46, with the same pattern seen in all NHP cortical layers, and in human STG: highest for delta, decreasing for theta, reaching a minimum for alpha, then increasing for beta and gamma. The ATRs add up to more than 1, greater than 100%, because of the oscillation overlap seen in **Fig 2**. Overlap was often due to nesting of a high frequency oscillation in a lower frequency event. For example, gamma and High gamma (Hgamma) both had high ATRs due to nesting in delta or theta bands. This pattern was observed both when using longer window sizes for slower oscillation frequencies compared to faster oscillation frequencies, and when using the full recording duration for the event rate calculations.

**Figure 6.**
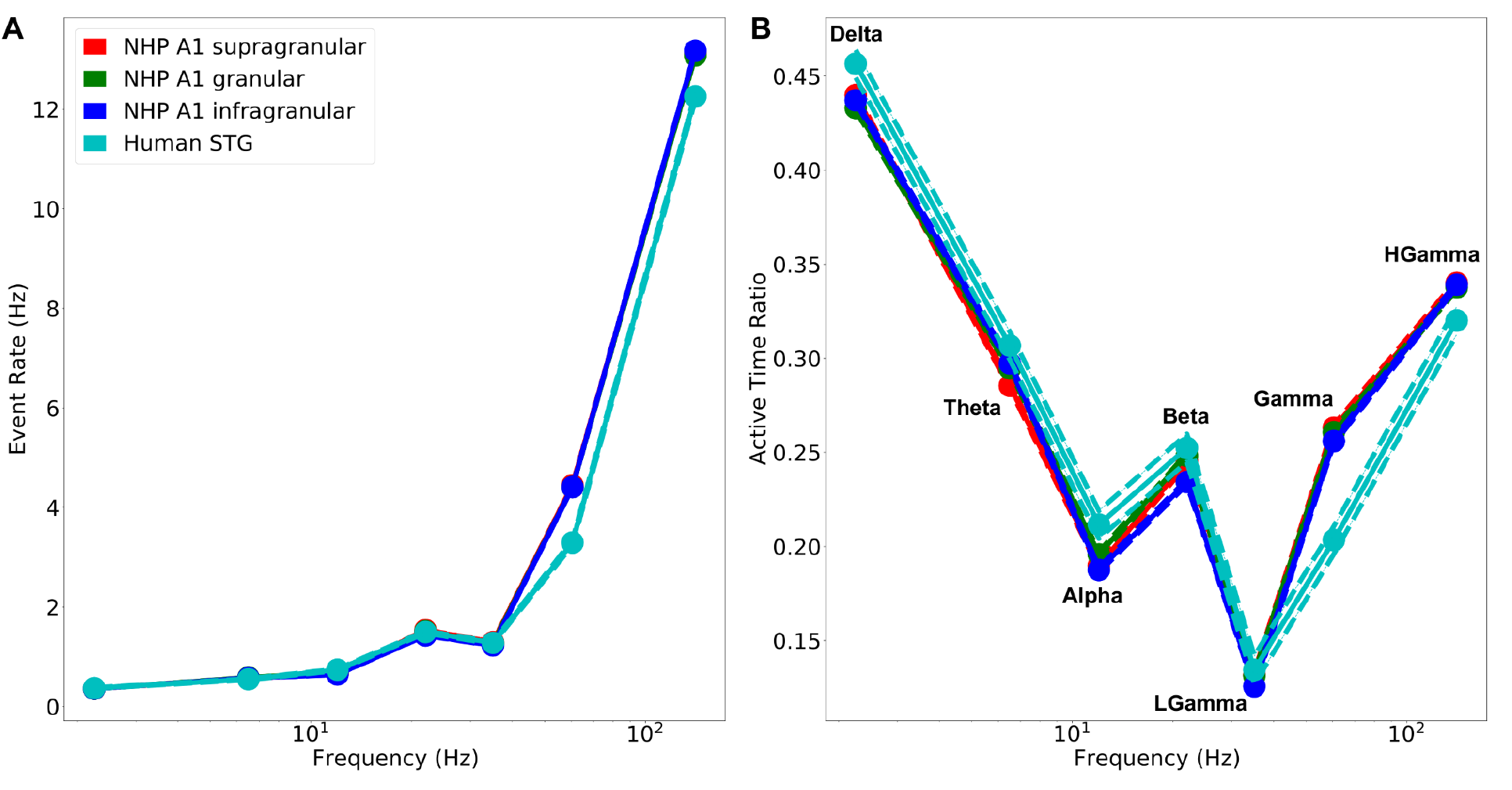
Oscillation event rates and active time varied by frequency band. **(A)** Higher frequency events are more frequent. (**B)** ATR: lower and higher frequency events fill much of the recording duration. (HGamma: high gamma bands; mean +/- SEM in both **A** and **B**).

While most oscillation characteristics were consistent across NHP layer locations, they were somewhat different when compared to human recordings (**Fig. 7**; **Supporting Material Tables 2-3**). The number of detected cycles averaged 3-4 (range 1-44), increasing from delta to high gamma (**Fig. 7A**). Time-domain count of local peaks closely matched the number of calculated cycles, demonstrating the accuracy of the measures taken in the wavelet domain (**Fig. 7B**). All oscillation frequency bands above delta had numerous events with >10 cycles in both NHP and human recordings. We defined intra-event frequency span as Fspan = log(maxF/minF) with minF,maxF the event’s minimum and maximum frequencies, respectively. Bandwidth was consistent across bands with intra-event Fspan showing maxF about 65% higher than minF (**Fig. 7C;** antilog(0.5)=1.65). A higher number of intra-event frequency shifts were seen in the delta band. Interestingly, human beta was broader in bandwidth, and human gamma tighter, compared to the values in NHP. Quality of filter-match increased with frequency in NHP (**Fig. 7D**). We believe that filter-match was worse at the lower frequencies due to overlapping higher frequency nested oscillations, that “distorted” quasi-sinusoidal wave shapes. This filter-match tendency differed in the human recordings, where high frequencies were poorly fitted by the filtered waveforms. The difference may be a technical consequence of more volume conducted activity, and thereby summation in iEEG recordings, but could also be a consequence of more intricate circuitry producing high frequency activity in humans vs. NHP. Future invasive recording/modeling studies will be needed to decipher this issue.

**Figure 7.**
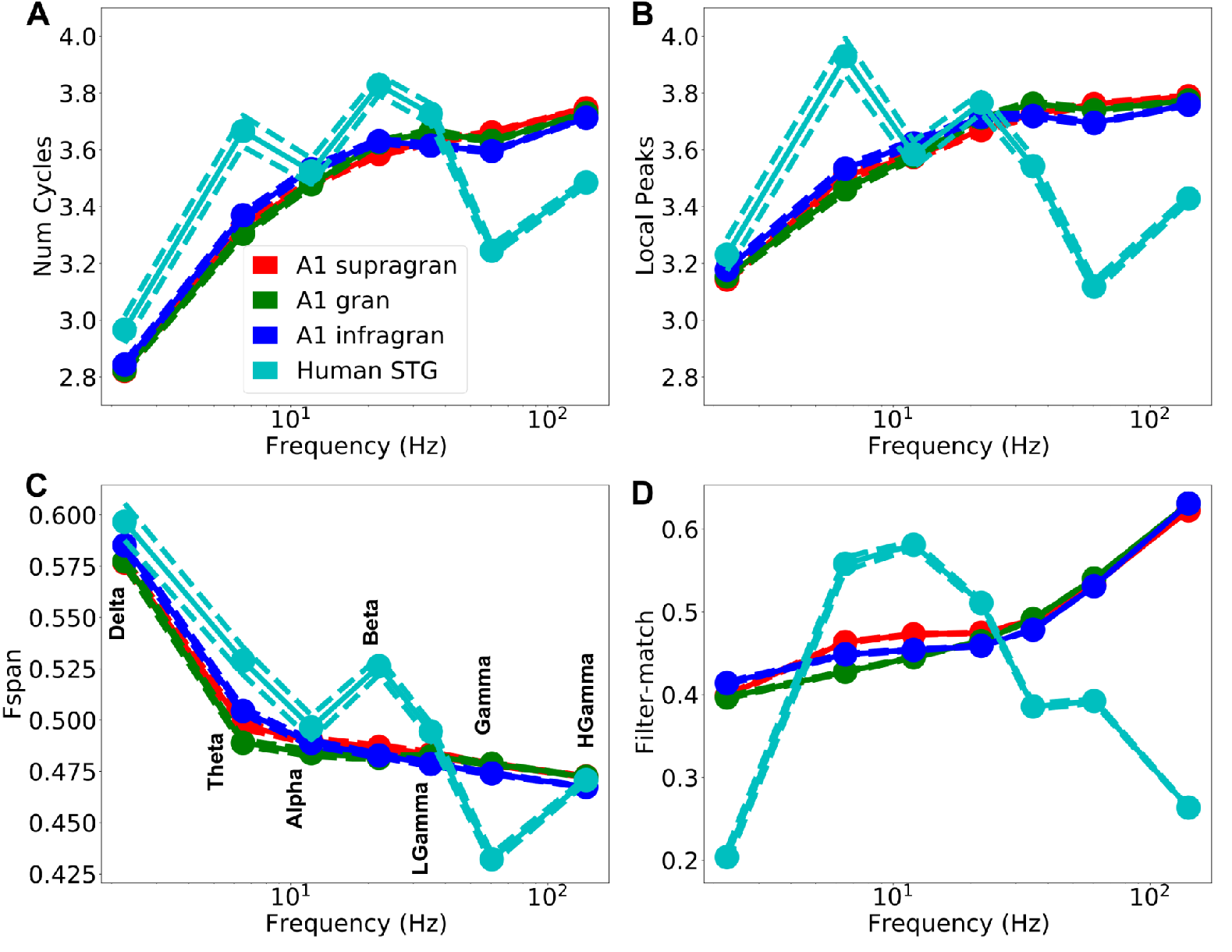
Event features. (A) Number of cycles. (B) Number of local peaks in the time domain of the filtered waveform. (C) Intra-event Fspan=log(maxF/minF); 0.7 is freq doubling. (D) filter-match *r* value.

A similar comparison was performed for multinuint activity (MUA) signals which tend to show higher number of cycles and lower filter-match values, but due to technical reasons (i.e. MUA is a noisier signal than LFP or iEEG due to environmental noise, movement, etc.), it is difficult to reliably interpret the observed differences between MUA and CSD characteristics from a physiological point of view. Nevertheless, the results of this analysis are shown in **Figure S6**, demonstrating that our methods can also be applied to the MUA signal.

Events occurred with some degree of rhythmicity, suggesting a rhythmic occurrence of these oscillation-events within windows of time (**Fig. 8**). The testing windows of 44.0, 30.0, 24.0, 10.7, 12.0, 3.6, 1.3 s for delta, theta, alpha, beta, low gamma, gamma, and high gamma were sufficient to allow about 16 events per window (see **Materials and Methods**). Inter-event intervals within a particular band were measured as times between event peaks, or by event initiation or termination (similar results). Coefficient of variation squared (CV2) and Fano-Factor (FF) were used as tests of rhythmicity. CV2 and FF are widely used measures in the literature of spike trains to describe their variability, with values intermediate between 0 (fully rhythmic and predictable) and 1 (fully noisy -- Poisson distributed). Additional details about these measures are provided in the **Materials and Methods** section.

**Figure 8.**
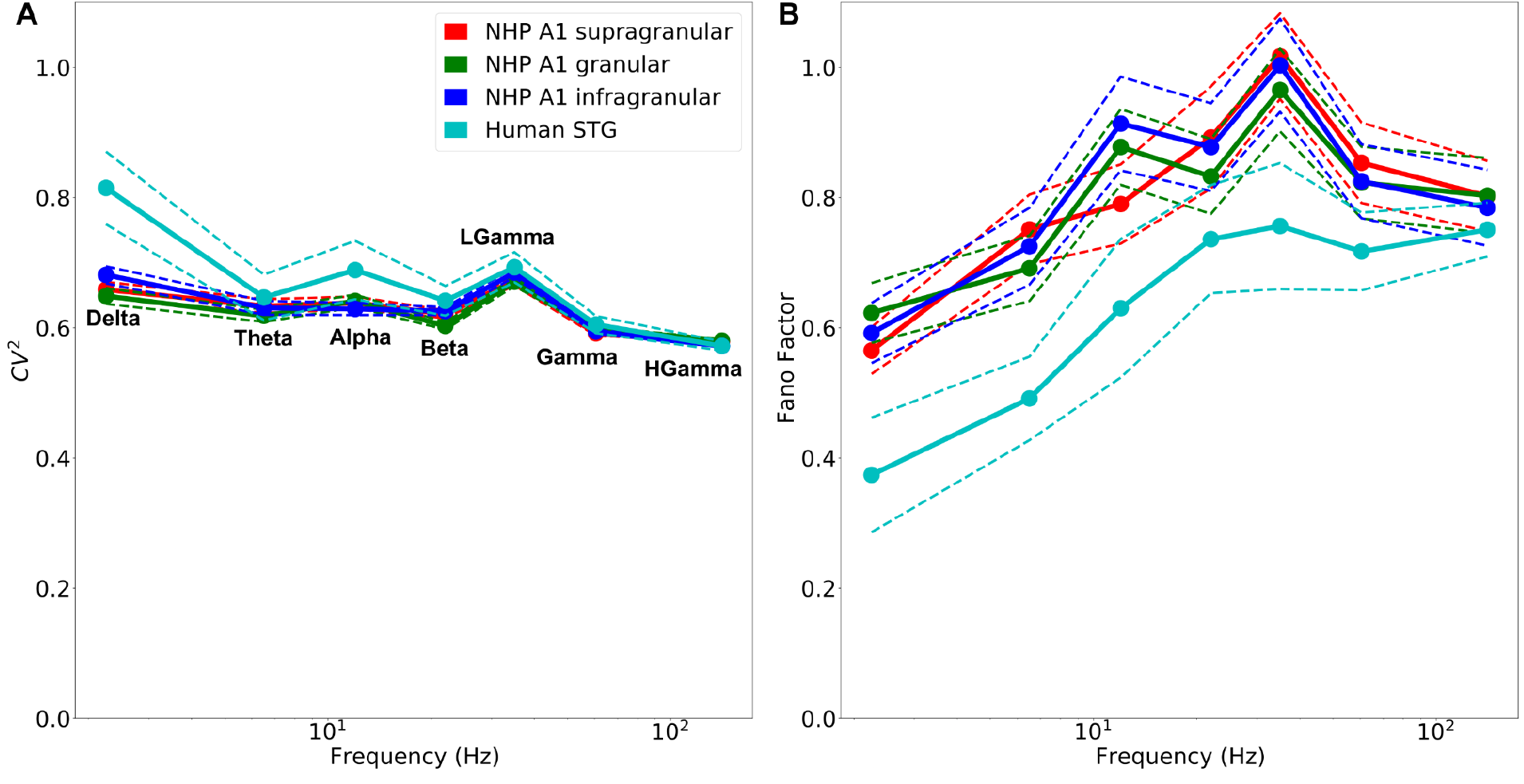
Inter-event intervals suggest rhythmic recurrence in all bands. **(A)** Squared coefficient of variation CV^2^ values (mean and SEM) in all bands < 1 (p<0.05, one-sided Wilcoxon signed-rank test).**(B)** Fano-Factor (mean and SEM) in all bands except for LGamma < 1 (p<0.05, one-sided Wilcoxon signed-rank test).

Average CV2 values were all substantially lower than 1.0 (**Fig. 8A**; p<0.05, one-sided Wilcoxon signed-rank test), demonstrating that events occurred across time with some rhythmicity. FF values were also consistent with this hypothesis, with mean values lower than 1.0 (**Fig. 8B;** p<0.05, one-sided Wilcoxon signed-rank test), with the exception of the LGamma band in NHPs, which had FF values slightly higher than 1.

### Oscillation events across bands

A common practice in the study of neuronal oscillations, which we adopted as well, is to divide oscillatory activity into specific bands of interest (i.e. delta, theta, alpha, beta, low gamma, gamma, high gamma). However, an oscillation event might span several of these bands. Thus, for each band of interest, we tested the percentage of band-limited events and the percentage of events that occupy multiple bands. To this end, we first defined the events based on the frequency at the peak amplitude of the event. We then tested whether the detected minimal and maximal frequency of each event (minF, maxF respectively) were both in the same band as the peak (band-limited), or if either crossed into other frequency bands. The results of this analysis are shown in **Fig. 9** for each band. Since all cortical layers showed similar results, we combined the data across layers in non-human primates. 15-25% of the events with a peak in the theta, beta, gamma and high gamma bands spread into lower frequencies, whereas approximately 40% of the events with a peak in the alpha range spread into the theta range as well. In addition, approximately 25-35% of events with a peak in the delta, theta, and gamma bands spread into higher frequencies, whereas 40-50%% of alpha and beta events spread into the beta and gamma band, respectively. In total, about 40-70% of delta, theta, gamma and high-gamma events were band-limited, while only 20-30% of events in alpha and beta bands were band-limited. While some of this variability across bands can be explained by the variability in bandwidth, this result indicates that some frequency bands tend to be more band-limited than others in A1 ongoing recordings. Note that in this analysis we used a combined gamma band (30-80 Hz) since low-gamma (30-40 Hz) is fairly narrow, which led to low-gamma having very few band-limited events with too many transitions from beta to low-gamma and from low-gamma to gamma (not shown).

**Figure 9.**
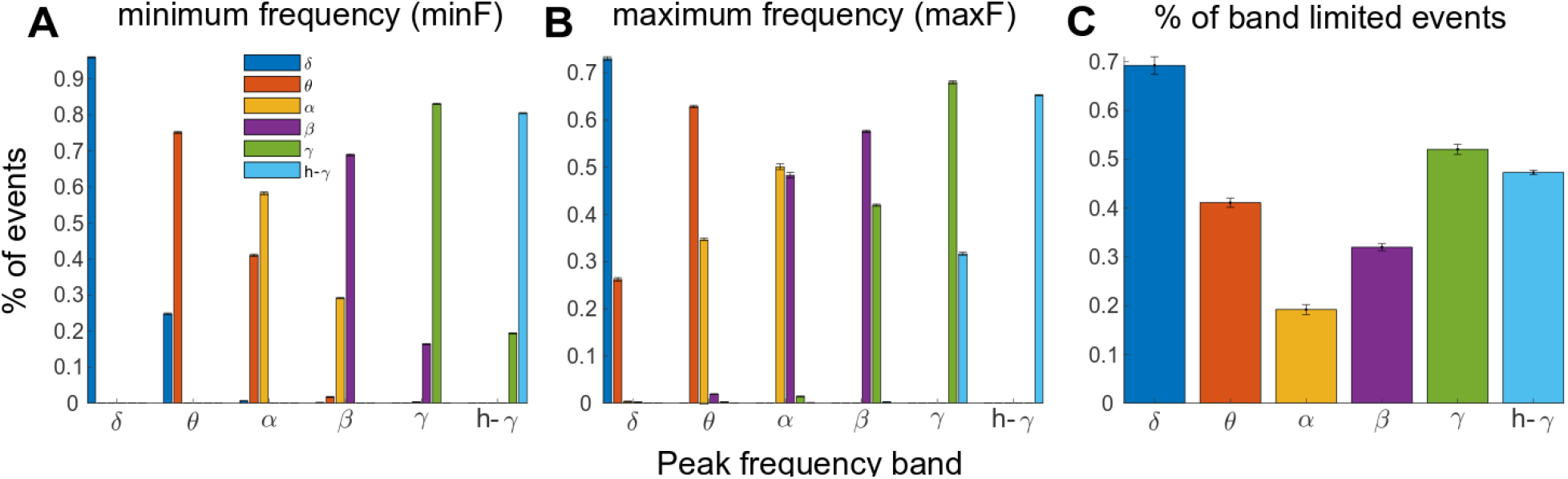
Individual oscillation events are categorized by the frequency of maximum amplitude (Peak frequency; x-axis) but may spread to lower or higher frequency bands. **(A)** and **(B)** display the % of oscillation events with minimum and maximum frequency in a specific band, while **(C)** displays the % of oscillation events that are band-limited, defined as having both minF and maxF in the same band as the event’s peak. In some frequency bands (i.e. delta, gamma and high gamma) individual events tend to be more band limited **(C)**, while in other bands (i.e. theta, alpha and beta) as many as 60%-80% of the events spread to adjacent bands. (X-axis location indicates categorization of an oscillation event based on its peak frequency; color indicates which frequency band an oscillation event spreads to - e.g. in **(A)** ∼75%,25% of theta events have a minimum frequency in the theta, delta bands respectively).

Neuronal oscillations have been suggested as foundational in auditory processing by ‘chunking’ the auditory stream into temporal units for further processing [29,30]. Specifically, oscillations in the delta-theta and gamma range seem to be engaged in tracking the dynamics of auditory information [31,32]. To further test the hypothesis that resting state auditory cortex dynamics are dominated by oscillations within these bands, we tested whether events in different frequency bands have higher probability of occurring together in the NHP auditory cortex. For each pair of frequency bands, we calculated the probability of co-occurrence as the number of events that occurred together (i.e. at the same time) out of the total number of events in these bands. Since the results were similar across the different cortical layers, we pooled the data from the different layers together. As predicted, the highest probability of events co-occurrence was found between delta-theta and gamma bands (**Fig. 10A**). Events in the alpha range had the lowest probability of co-occurrence with any other frequency band. These results indicate that transient oscillation events within the delta-theta and gamma frequency bands are inter-connected in the auditory cortex even in the absence of any auditory stimuli. This might indicate distinct delta-theta vs. alpha-beta dominated brain states, as one of our former [16] and another study demonstrated [32].

**Figure 10.**
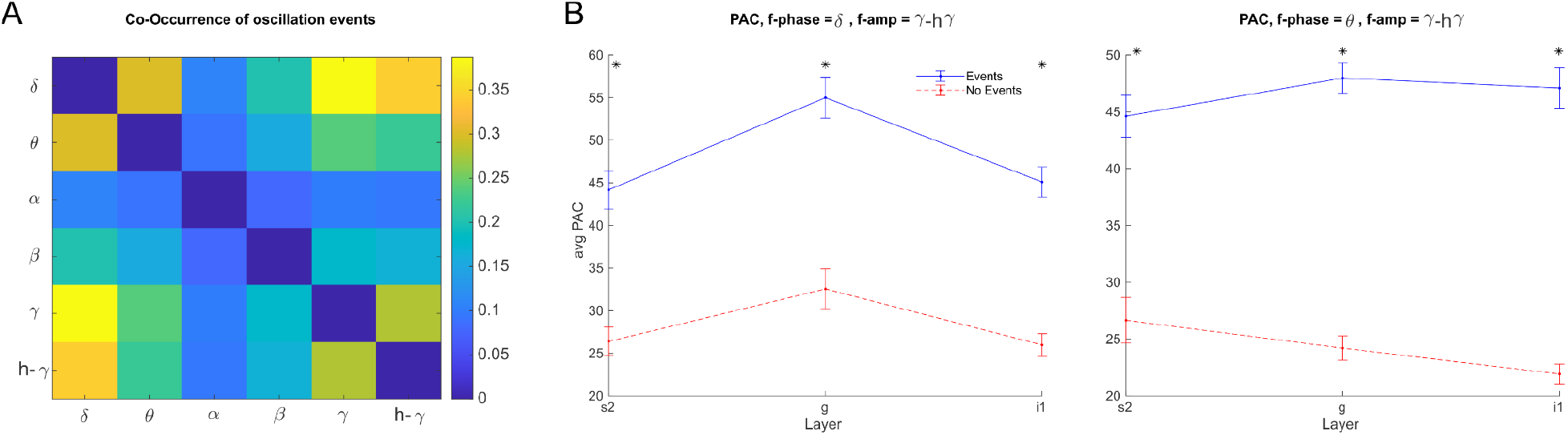
Co-occurrence of oscillation events and phase-amplitude coupling. **(A)** Oscillation events in the delta-theta range had higher probability of occurring together with gamma-high gamma events. **(B)** PAC between delta (left) or theta (right) events and gamma/h-gamma events during periods of low-frequency events (blue) and periods without any detected events (red). PAC in all layers increases during periods of low-frequency events. (Note that just as in the analysis shown in **Fig. 9**, here we used a gamma band spanning 30-80 Hz, to allow accurate measurement of cross-frequency interactions in a wide enough band).

To further test the interaction between delta-theta and gamma oscillations, we calculated the phase-amplitude coupling (PAC) between these bands during delta and theta oscillation events and compared them to segments without oscillation events from the same recordings. Both delta and theta bands showed an increase in PAC with gamma band oscillations during the oscillation events compared with no-event segments (t-test; p < 0.05, Bonferroni corrected), indicating that gamma events tend to appear at specific phases during low-frequency oscillations (**Fig. 10B**).

## Discussion

The current study was aimed at characterizing basic features of individual oscillation events in the auditory cortex of NHPs and humans. These features and the variability across events both within and across frequency bands demonstrates the importance of studying singular oscillation events, as opposed to averaging them, as each event might contain information related to its functional role. Studying single events will also aid in revealing their underlying mechanisms and deficits in neuropsychiatric disorders.

We found near-continuous (90% of time) neuronal oscillations across the full range of frequency bands in a large, resting-state electrophysiology dataset invasively recorded from non-human primates and humans. Oscillations occurred in *events* of up to 44 cycles (**Fig 5F; Supporting Material Table 2**), with about 3-4 cycles on average (**Fig. 7A**). We developed a software package, *OEvent*, that used a wavelet transform to identify and characterize these events from CSD and iEEG signals, after removal of event-related potentials and broadband noise. OEvent was validated using simulated oscillations at different frequency bands that were superimposed on the fluctuating background from NHP recordings (**Fig. 1**).

We determined basic features of oscillation events: power, frequency at peak power, duration, frequency span (bandwidth), number of cycles, and filter-match (quantifies presence of pure oscillation) (**Fig. 7D**). The frequency of oscillation events remained within a single traditional frequency-band, but it was far from being constant in most cases within the event (*e*.*g*., U-shape in **Fig. 3,4** alpha). Intra-event shift was on average about 65% (indicated by Fspan of ∼0.5 in **Fig. 7C**). Interestingly, oscillation events in a particular band occurred with some regularity, demonstrated by noting CV2 and Fano Factor values significantly less than 1. This was likely due to cross-frequency coupling effects. Overall, our results provide evidence that neuronal oscillations across a wide range of frequencies occur in quasi-rhythmic, multicycle events, or “bursts”, with significant across-burst temporal predictability. We suggest that rhythmic oscillatory bursting is the dominant operational mode of the auditory cortex in both human and non-human primates.

Acoustic signals usually contain information at multiple time scales, distinguishing between rapid changes and the general envelope of the signal [33]. As shown by our ATR analysis, in both humans and NHPs, delta and gamma/high gamma oscillations exhibit the highest ATR. Moreover, PAC between delta/theta and gamma/high gamma oscillations is increased during oscillation events (**Fig. 10**). In humans, these bands correspond to the rates of phonemes and syllables in speech [34,35]. Non-task related oscillatory bursting at these specific bands might serve to maintain temporal coupling of neuronal activity and indicate the ‘readiness’ of the auditory cortex to process auditory information containing acoustic modulations at multiple time scales.

### Oscillation event variability

Several previous studies have used rigid criteria to identify oscillations, often only identifying an oscillation if it showed a single frequency, and relatively constant amplitude, across multiple cycles [36,37]. We have opted to cast our net more widely in order to capture the full variety of oscillatory phenomenology. Our main reason for this broad catchment is that we see major frequency and power shifts in a context where oscillations have a clear functional implication - for example in the delta/theta/alpha gamma modulations seen during auditory processing [31,32]. Strict criteria also do not adequately take into account the nonstationarity (formerly called noisiness) of brain signals due to constantly fluctuating cognitive and behavioral demands unrelated to task [38]. By taking a less restrictive approach, we discovered events with variation in cycle-to-cycle frequency and amplitude. Admittedly, the detection/characterization of oscillations will likely always remain a signal to noise issue, but we believe that by attaching the properties to oscillation events described here, we can get closer to detecting “real oscillations” in the brain.

Interpretation of rhythms, and an establishment of identities for conserved rhythmic “motifs” relies first on measurement technology and then on methods of analysis [39]. Intracranial and extracranial recordings produce vastly different signals for interpretation that may require different tools. Even within intracranial recordings, there are important differences between depth and surface recordings, between monopolar vs bipolar recordings, and between LFP and CSD. As shown by our simulations, different frequency bands and even different numbers of cycles within a band may also require different approaches, particularly in cases of unusual stereotyped events such as ripples. Different brain areas and different task conditions may also provide different signal patterns to be taken into consideration when identifying oscillation events. In the current study, we were encouraged by obtaining similar results across different measurement technologies (iEEG, LFP, CSD, MUA) and across species in resting state conditions.

In the data we analyzed, there were intuitively clear examples of oscillations with high power, multiple cycles, relatively constrained frequency range, and sinusoidal appearance (**Fig. 2-5**), features typically considered when determining whether a portion of a signal contains an oscillation. Individuals will nonetheless disagree on the exact cut-offs to make this “oscillatory determination”. This subjectivity makes it difficult for the neuroscience community to arrive at a consensus on how to determine the importance of oscillations in the brain. A further complication is that data used in publications is rarely presented in “raw form.” We have therefore presented a heterogeneous portion of the dataset used in the Results in **Fig. 5**, to allow readers to view examples of oscillations with all of their variability. The presentation of raw data can provide confidence that the oscillations under scrutiny are not simply “filter artifacts”. For example, since varying wavelet filter properties could potentially lead to different values for the number of detected cycles in an oscillation, researchers should carefully examine their raw data and make sure the methods are appropriate for their particular research goals and questions. For these reasons, we recommend sharing raw data for all studies involving oscillations.

The contrast between our identification of high oscillation variability with a more rigid interpretation of oscillation identity can be seen by comparing our techniques with those of Voytek and colleagues [36], which only considered oscillations with consistent features over time. Their method first performed low-pass filtering to remove high-frequency noise, followed by band-pass filtering to narrow-in on a frequency of interest. Next, their algorithm extracted local minima/maxima of individual cycles in the filtered time-domain signal, and analyzed cycle-by-cycle properties, checking for consistency of features over time [20,36]. Voytek’s algorithm thereby provided criteria for defining stationary neural oscillations by assuming consistency of cycle-to-cycle amplitude and frequency during the single event. The algorithm also used a minimum threshold for the number of cycles in an oscillation. While this approach is very useful, we argue that at first, we should detect all potential oscillation events, extract their basic properties, and then clearly state which properties identify actual oscillation events. Of course these criteria will be subjective for a while for each research group, but if we agree on the main measures, this will make any oscillation based result more replicable.

In the future, collaborative software tools and additional data sharing will improve consistency in the research community and make faster progress. To this end, we are sharing the current datasets. Researchers can use our datasets and accompanying OEvent software package (https://github.com/NathanKlineInstitute/OEvent) to rate rhythmicity within and among events, to evaluate and compare event detection tools, and to develop new tools. Further progress would come from embedding software platforms in web front-ends to allow researchers to view neurophysiological data and perform oscillation scoring on signals in public datasets or from their own data. This process would help the community come to a consensus on neurophysiological oscillations and the features that contribute to oscillations *vs*. event-like waveforms, as well as objectively defining rhythmicity using measures such as coefficient of variation squared (CV2), Fano Factor, or a modified version of lagged coherence that operates on events [40]. Consistent consensus across studies may be somewhat limited by different signal patterns in different brain areas or under different task conditions.

Other published methods have analyzed signals in only the frequency domain [41] or in only the time domain [42]. As shown in the present paper, there are clear advantages in moving back and forth across these different views into dynamics. Here, we were able to more accurately exclude ERPs using features from both domains: *e*.*g*., waveform shape and duration (time-domain), along with frequency-span (frequency-domain). Our analysis on the quasi-rhythmicity of oscillation events relied on first defining the events using the frequency domain (wavelet transform) and then using time-domain to measure inter-event rhythmicity (CV2 of interevent intervals).

Using the methodology presented here and also methods from other labs, our plan is to determine signatures or motifs of particular oscillation types in corresponding locations in human and non-human primates. This will consist of looking at patterns of crescendos and decrescendos in activation power, or patterns of increases and decreases in frequency. For example in the alpha trace shown in **Fig. 3**, the wavelet pattern shows a rapid increase in power and then gradual decrease, while the frequency first decreases and then increases. We also plan to incorporate methods that are commonly used for spike sorting to distinguish between different vs. repeating waveforms of oscillatory activity, which might shed light on their underlying physiological mechanisms.

### Mechanisms of oscillation generation

Biophysical computer modeling of neurons [43,44] and detailed microcircuits can also help define oscillations and their properties, by allowing predictions about the mechanistic origins and temporal fluctuations of specific oscillation types and their recurrence over time [45,46], and how different oscillation generators interact [25,26,47–49].

Detailed computational modeling of neural circuits has shown how interactions between different classes of interneurons contribute to fast and slow rhythms, via short and long GABA_A_ synaptic time constants [50–53]. *In vivo*, we expect that interactions between different interneuron sub-populations and pyramidal neurons will result in a mix of oscillations in the recorded signals [54–58], making it unlikely to observe a constant frequency/amplitude oscillation. Some of these interactions, which could lead to shifts in peak oscillation frequency over time or variability in waveform shapes in different nearly simultaneous oscillation events, could be explored through detailed computational modeling. In addition, differences in recording methods (electrode properties and volume recorded from) between NHP and humans could be investigated with detailed biophysical modeling.

Data-driven modeling could also pave the way for understanding the extent to which electrophysiological measurements are contaminated by noise, and which type of waveforms one can expect to see in electrophysiological signals recorded during different experimental conditions. Biophysical modeling could further allow us to develop a revised taxonomy of the different oscillations, by predicting which circuit components each oscillation arises from [48,59–61]. This type of modeling can be taken a step further by delineating how circuit features contribute to oscillation event dynamics that support particular aspects of auditory processing, such as speech tracking [62–64] or auditory steady-state responses [65–67]. A clearer understanding of the mechanisms creating oscillations will also pave the way to improving neuropathologies associated with disrupted brain rhythms, via pharmacology or targeted neuromodulation that normalizes rhythmic activity in a principled manner [68].

### Ubiquity of oscillation events

To our knowledge, ours is the first study to systematically quantify the full range of physiological oscillations in the auditory cortex, using the novel method of oscillation event analysis. Prior studies have shown that high-power beta rhythms emerge as brief transient events lasting only a few cycles, in somatosensory [21,24], motor [69–71] and frontal cortex [21,72]. In somatosensory cortex, extracranial measurement in humans, and intracranial in mice and NHP [21,24] showed a low rate of beta events comparable to what we observed (1.4-1.5 Hz; **Supporting Material Table 1A**). We found 3.6-3.8 cycles per event (**Supporting Material Table 2**) somewhat higher than the less than three cycles found in the prior studies. Our merging of events when bounding boxes overlapped by 50%, could produce our somewhat larger number of cycles. In addition, in our present study we used a lower threshold (4x median compared to the 6x median threshold used previously), and thereby potentially allowed detection of a larger set of oscillation events with different properties. Further research is required to determine if there are more similarities between beta bursts in the auditory cortex as compared to other brain areas and studies.

Consistent with our analyses, a study on cat auditory cortex neuronal activity demonstrated that gamma rhythms are non-stationary, and event-like [22]. The study also found that gamma events’ peak frequency was dependent on arousal level, while amplitude was more dependent on attention. A study on human auditory cortex found precise and distinct timing to the onset and offset of beta vs gamma oscillations during musical beat processing [73], suggesting different auditory processing roles for the two frequency bands. Related research in human auditory cortex supports the role of event-like beta and gamma oscillations in auditory processing and auditory-motor interactions [74–77]. Other studies show that both beta and gamma bursts together contribute to working memory processes [23]. These, and our current results highlight the importance of characterizing multiple “oscillatory dimensions”, in order to better understand how different oscillatory features relate to specific brain functions.

Although these studies did not specifically quantify predictability of gamma recurrence over time during spontaneous activity, related evidence from the auditory cortex [78] and other brain areas [79] shows that the phase of low-frequency rhythms (delta/theta/alpha) influences the amplitude of higher-frequency rhythms (beta, gamma) in a predictable way, which is termed phase amplitude coupling [80–82]. Other recent studies show that apart from predictability in time, there is predictability in the spatial location where gamma bursts occur: precisely orchestrated alpha waves traveling over the cortical surface regulate the timing of bursts of localized gamma activity [83]. Another recent study proposed a promising method, called superlet, that uses sets of wavelets that are combined geometrically to maintain the good temporal resolution of single wavelets and gain frequency resolution in upper bands (Moca et al., 2021).

### Functional implications

While our current study was specifically designed to be non-task-related, no effort was made to make the subjects cognitively or perceptually inactive. Although the term *resting-state* is applied to this condition, the brain is never functionally at rest [38,84,85]. We argue that “resting state”, at least for our study, provides the most colorful palette of ever changing brain activity, and therefore, the most variation in oscillation events. The brain activity we have measured is that of a perceiving, thinking brain -- hence we prefer *non-task-related state* to *resting state*.

Oscillations are sometimes considered to be functionally irrelevant, or epiphenomenological. By contrast, we and others view oscillations as fundamental to brain function and brain encodings, and view understanding the roles of oscillations as being critical for understanding the brain [86]. In a trivial sense, oscillations are epiphenomena since they reflect an electric field that is only an external reflection of internal dynamics [87]; in this same sense an external recording of an action potential is an epiphenomenon. The important question is whether aspects of the oscillation, or aspects of a single-neuron spike train or aspects of oxygenation state in a brain area, can be regarded as having a role in coding and representations [88,89]. The notion of coding itself then breaks into 2 meanings: correlative coding that can be observed by the experimenter vs. causal, internal coding that forms part of how the animal behaves or thinks. Correlative coding is more studied; causal coding is more important.

As with the single-neuron firing-rate causal coding hypothesis, evidence for oscillation causal coding is inferential and hypothetical. The major hypotheses on the role of oscillations are entrainment theory [5–7] and communication through coherence theory [9,12]. Grossly, both of these theories consider the role of phase synchronization in gating communication channels across brain circuits. Specific frequency bands have been implicated in particular aspects of neural state: attention indexing by alpha in auditory, visual and somatosensory areas [15,16,90,91]; stimulus timing by aligned delta-theta [92,93]; delta with nested gamma by encoding of elements of speech signals [31,94,95]. The present study demonstrates that oscillations are ubiquitous across bands and thus would provide fertile ground for their use in these contexts. At a fundamental level, the oscillations we observed reflect ensembles of neurons activated synchronously and projecting activity together both within and between areas.

An additional hypothesized oscillation role involves the control and molding of higher frequency oscillation nested within lower [96]. Evidence for such frequency nesting was seen by our filter-match measure (**Fig 7D**), which measured the difference between a clean, but not necessarily single frequency, sinusoid and the recorded wave and by co-occurrence and PAC analysis (**Fig. 10**), which shows an increase in the probability of events occuring at the same time and during a certain phase for specific bands. Further work remains to be done in determining the ranges and relations of particular nested frequencies relative to particular nesting frequency. We also observed another indication of oscillation variability, the ∼65% variation in frequency of a primary oscillation, demonstrated by the Fspan measure in many events (**Fig 7C**). Both nested oscillations and oscillation frequency-spread could be features of entrainment theory and communication through coherence. Alternatively, one or both of these might be an indicator of an underlying embedded encoding.

In conclusion, we argue that the oscillatory features extracted in the current study provide a richer view on oscillatory brain activity. The suggested features can be further explored in a task-related context and in neurological disorders to better understand the role of neuronal oscillations in information processing and in disease [97–100].

## Materials and Methods

### Datasets

We used two datasets of neuronal activity, invasively recorded over longer time scales (minutes) in non-task related conditions: 1) laminar electrode array local field potentials from non-human primate (NHP) primary auditory cortex (A1; 92 recordings totaling 487.8 minutes); 2) intracranial EEG (iEEG) from human superior temporal gyrus (STG; 8 recordings totaling 37 minutes). In all analyses (except for specific examples), the data was pooled across subjects. However, examination of individuals showed minimal variability for most oscillation event features (individual summaries not shown).

1. NHP electrophysiological data was recorded during acute penetrations of area A1 of the auditory cortex of 1 male and 3 female rhesus macaques weighing 5-8 kg, who had been prepared surgically for chronic awake electrophysiological recordings. Prior to surgery, each animal was adapted to a custom fitted primate chair and to the recording chamber. All procedures were approved in advance by the Animal Care and Use Committee of the Nathan Kline Institute. Preparation of subjects for chronic awake intracortical recording was performed under general anesthesia using aseptic techniques. To provide access to the brain, either Cilux (Crist Instruments) or Polyetheretherketone (PEEK; Rogue Research Inc.) recording chambers were positioned normal to the cortical surface of the superior temporal plane for orthogonal penetration of A1. These recording chambers and either socketed Plexiglass bars or a PEEK headpost (both used to permit painless head restraint) were secured to the skull with ceramic screws and embedded in dental acrylic. Each NHP was given a minimum of 6 weeks for post-operative recovery before behavioral training and data collection began. Before and after recordings, NHPs had full access to fluids and food. No rewards were offered during recordings. The data were recorded during waking rest with eyes mostly open, while the macaques were in a dark room. During recordings, NHPs were head-fixed and linear array multielectrodes (23 contacts with 100, 125 or 150µm intercontact spacing, Plexon Inc.) were acutely positioned to sample all cortical layers of A1. Neuroelectric signals were continuously recorded with a sampling rate of 44 kHz using the Alpha Omega SnR system. We note that since the monkeys were in a dark room, we would not expect visual inputs to influence the outcome in the auditory cortex. However, we know that eye movements influence oscillatory activity in auditory cortex even in the dark, and based on some recent results from the lab (unpublished), eyes closed vs eyes open might influence dynamics in the auditory cortex as well.
2. The iEEG data, previously published [101,102], was recorded at Northshore University Hospital in medically-intractable epilepsy patients that underwent stereotactic depth electrode placement as part of their epilepsy surgery workup. Electrode placement was based on clinical criteria only. All patients provided written informed consent monitored by the institutional review board at the Feinstein Institutes for Medical Research. Electrode localization methods have been described previously [103]. The data was recorded using a Tucker Davis Technologies amplifier with a sampling rate of 3000Hz. The data were recorded during waking rest with eyes closed. While we cannot exclude a possible effect of medications on brain oscillations in general, a systematic effect is unlikely because: 1) these recordings are usually done when patients are either off medications or being tapered off, 2) different patients are administered multiple different medications which makes a systematic effect across patients unlikely, and 3) the patients were not administered barbiturates. Benzodiazepines were only used to abort seizures and we avoided recording experiments following seizures that require its administration. Note that our recording systems store the iEEG data in either V or µV formats, depending on the recording system used. **Fig. 4** values are in Volts.

### Data processing

All analyses from the NHP dataset were run on current-source density (CSD) signals, calculated as the second spatial derivative of laminar local field potential recordings. This was done to reduce potential issues related to volume conducted activity. We estimate CSD using laminar local field potential recordings. To reduce computation time, we only used channels that included the supragranular, granular, and infra-granular current sinks in response to preferred frequency tones, considered “active” since they measure depolarizing transmembrane currents [16]. In humans, it is not as simple to calculate CSD due to more restricted electrode arrangements. Thus, all analyses from the iEEG dataset used the recorded signals themselves without taking any spatial derivative (re-referencing signals using a bipolar reference produced similar results; not shown).

### Removal of externally-driven events

Prior to event detection and feature extraction, we removed signals that are suspected to be the results of Event-related potentials (ERPs). ERPs are prominent brief signals associated with external sensory stimuli. The wavelet transform might identify these transient signals as oscillations, obscuring our analysis. In order to remove these, we formed average ERP waveforms in supragranular, granular, and infragranular sink channels from an NHP A1 dataset, recorded during 50 dB auditory click stimulus presentations (**Fig. 12**). The wavelet transforms of these signals showed high spectral power. Both the supragranular and infragranular ERP responses have sharp peaks that last around 50 ms, producing a 20 Hz signal in the wavelet analysis. Similarly, the granular ERP response has a slower component that lasts for approximately 100 ms, producing a 10 Hz response in the wavelet spectrogram.

**Figure 11.**
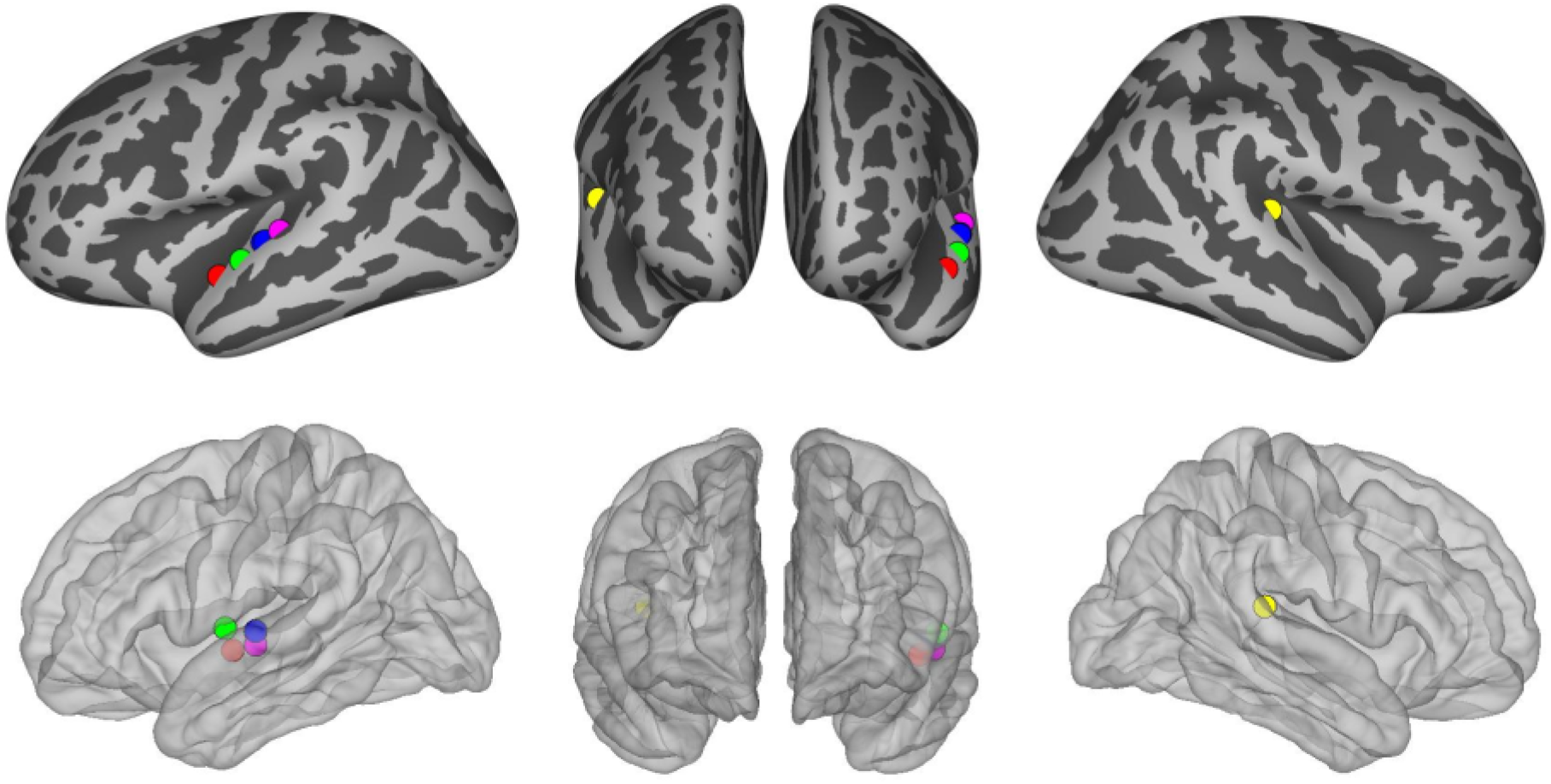
Locations of the intracranial EEG electrodes used for human electrophysiology recordings included in this study, overlayed on a standard average brain. Colors represent different patients.

**Figure 12.**
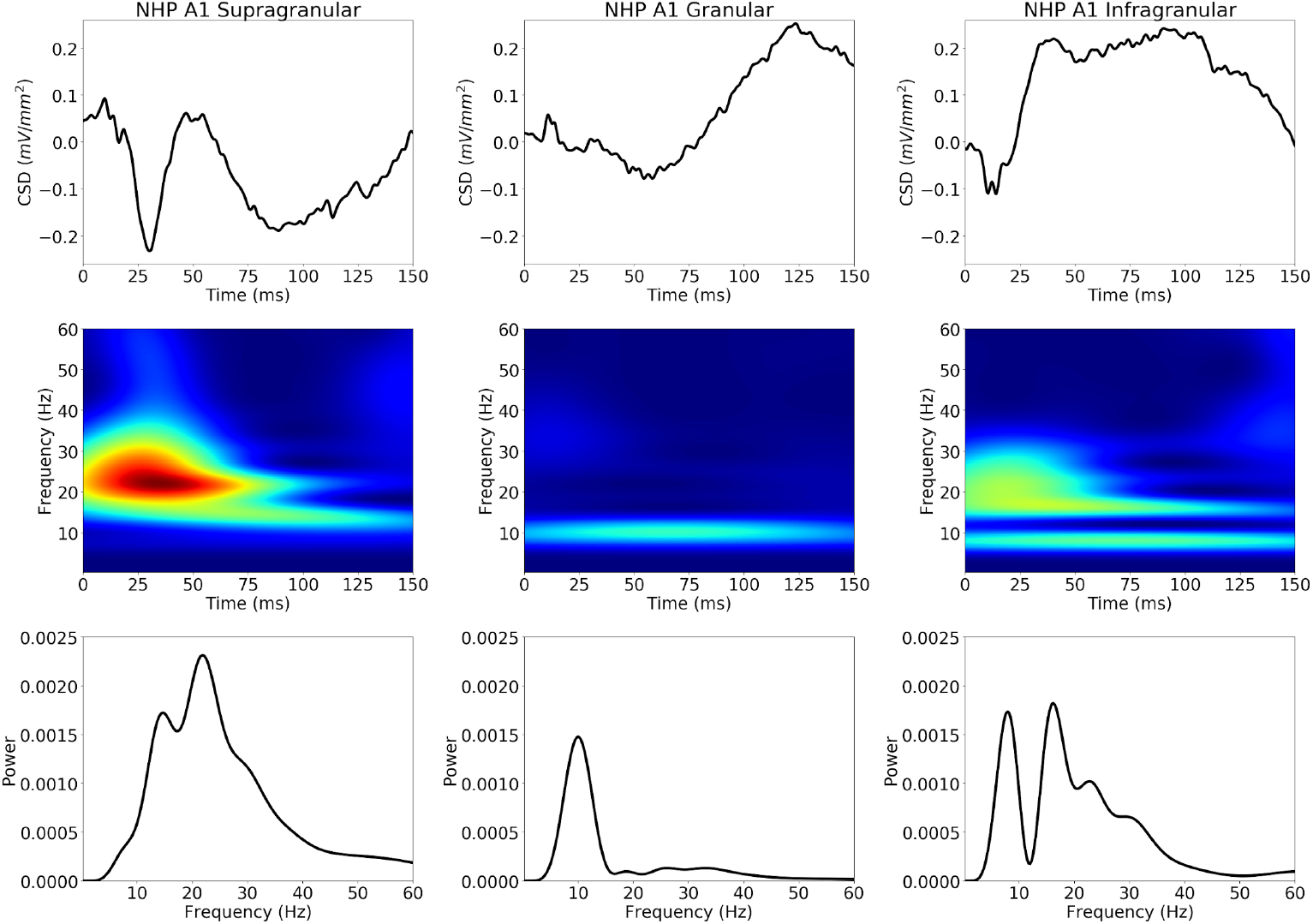
Stereotyped ERPs in NHP A1: supragranular, granular, infragranular layers (left to right; 50 dB clicks in NHP). **Top:** Average ERP waveforms (click at 0 ms). **Middle:** wavelet transform spectrograms. **Bottom:** Apparent oscillation peaks (average of spectrogram over time).

ERP score was defined as the maximum normalized cross-correlation between that event and the average ERP from the same cortical layer (**Fig. 12, bottom**). An event with ERP score >0.8 with duration 75-300 ms (same range used for all layers; a loose constraint to reduce false negatives) was excluded from further analysis. ERP score was calculated for all NHP A1 events, but not for human STG where we lacked ERP data. We also excluded events with broadband frequency responses, likely from rapid-onset ERPs or recording noise. We defined logarithmic frequency span as Fspan = log(maxF/minF) using natural log, and excluded events with Fspan>1.5 (2.17 octaves).

### Oscillation event (OEvent) detection and feature extraction

We extracted moderate/high-power spectral events using 7-cycle Morlet wavelets on non-overlapping 10 s windows [21,49]. The 7-cycle Morlet wavelets were chosen to provide an adequate compromise between time and frequency resolution [18]. We used linearly spaced frequencies (0.25 Hz frequency increments), ranging from 0.25 - 250 Hz, to compute the wavelets. The power time-series of each wavelet transform was normalized by median power across the full recording duration. We then applied a local maximum filter to detect peaks in the wavelet transform spectrogram. All local peaks were assessed to determine whether their power value exceeded a threshold. We used a moderate power threshold (4× median) to determine the occurrence of moderate-to high-power events.

A local power peak in the spectrogram was defined within the local 3×3 window it was centered in, and exceeded the 4× median threshold of each individual frequency. The frequency and time bounds around that peak were determined by including time and frequency values before/after and above/below the peak frequency until the power value fell below the smaller of ½ maximum event amplitude and 4× median threshold. As shown in **Fig. 2**, this produces a bounding box around each oscillation event that can be used to determine frequency spread (minF to maxF), time span (start, stop), and peak frequency (defined as the frequency at which maximum wavelet power is detected). After the initial set of oscillation events is detected, we merge events when their bounding box overlapping area in the wavelet spectrogram exceeds 50% of the minimum area of each individual event. This allows continuity of events that are separated by minor fluctuations below threshold.

We then calculated additional features from this set of events. We calculated the number of cycles by multiplying the event duration by its peak frequency. We also filtered the underlying signals of each event using a zero-phase shift bandpass filter within the minF and maxF frequency ranges, defined on a per-event basis (as in bounding boxes shown in **Fig. 2**). We calculated filter-match value *r*, defined as the Pearson correlation between this filtered signal and the raw signal, and used it as an index of how clearly the event oscillation is visible in the raw signal. Based on visual inspection of numerous waveforms, we suggest associating the following ranges of filter-match values with the corresponding qualitative assessments (0.0-0.25:weak; 0.25-0.5:moderate; >0.5:strong/high). Using the filtered signal also allowed us to count other features of the oscillation, including number of peaks (local maxima) and number of troughs (local minima). Number of peaks and the number of cycles were highly correlated. In figures showing waveforms of individual events, the 0 time alignment is taken as the wavelet transform 0 phase closest to the time of event threshold.

After extracting individual oscillation events, we classified them into the standard physiological oscillation frequency bands on the following intervals: delta (0.5-4 Hz), theta (4-9 Hz), alpha (9-15 Hz), beta (15-29 Hz), low gamma (30-40 Hz), gamma (40-80 Hz), high gamma (81-200 Hz). This classification was based on the frequency at which maximum power occurred during each event (intervals were open on the lower bound and closed on the upper bound).

We compared our OEvent method with the BOSC algorithm [27] as shown in **Fig. 1**.

### Phase-amplitude coupling calculations

Phase-amplitude coupling was calculated by filtering the entire recording via convolution with a complex Morlet wavelet of width 3 and extracting the instantaneous phase and amplitude at the appropriate frequencies (f-phase = delta - 0.5-4 Hz; theta – 4-8 Hz; f-amplitude = gamma – 30 – 200 Hz). Then, the instantaneous phase and amplitude were segmented based on the timing of the low-frequency events. Modulation index for segments containing low-frequency events and segments without any detected events was calculated (matched for number of segment and segment duration) using: *z*(*t*) = *Amp*_*r*_(*t*) *e*_*δ*/θ_^*iφ*(*t*)^. The modulation index is defined as the mean of *z*(*t*) and provides a metric of coupling between the two time series. We then averaged the modulation index across segments with or without low-frequency events and compared the averages using a t-test.

### Inter-event interval characterization

To measure rhythmicity across events from a given oscillation frequency band, we formed inter-event interval (IEI) distributions from the oscillation events which were not characterized as ERPs (see **Results**). We formed the intervals in two ways: 1) the interval between the time of peak power of the previous event to the time of peak power of the next event; 2) the interval between the end of the previous event to the start of the next event. After forming IEI distributions, we calculated its squared coefficient of variation (CV2). We also calculated the Fano Factor (FF) from the number of events in successive windows (defined below). The FF is defined as the variance to mean ratio of a random process in a certain time window, while the CV2 is defined as the variance over the squared mean. Both measures describe the extent of variability in relation to the mean. A Poisson distribution will have CV2 and FF values of 1, a more rhythmic process will have CV2 < 1, while a “bursty” process (multiple characteristic inter-event intervals) will have CV2 > 1. CV2 values increased with the number of events in a time window, for both longer windows of analysis and higher frequency oscillations. To control for this, we varied window size for different frequencies (generally longer for slower frequencies) to produce a similar number of events per window (N ranging between 12-18, depending on frequency band and species). The empirically-determined window sizes used were 44.0, 30.0, 24.0, 10.7, 12.0, 3.6, 1.3 s for delta, theta, alpha, beta, low gamma, gamma, and high gamma frequency oscillations, respectively. Note that the window size for low gamma events was longer than for beta beta events, since low gamma events occurred less often. We used a one-sided Wilcoxon signed-rank test to determine that the measured average CV2 and FF values were lower than those of a Poisson process.

### Source code

Python source code for our OEvent software package, for oscillation event detection and analysis is available on github at https://github.com/NathanKlineInstitute/OEvent

## Acknowledgments

Research supported by NIH R01DC012947, Army Research Office W911NF-19-1-0402, New York State ECRIP Fellowship, NIH P50 MH109429, NIH R01MH106174, NIH U24EB028998, NIH U01EB017695, NYS SCIRB DOH01-C32250GG-3450000, NSF 1904444, NIH R01MH111439, and a grant from The James S. McDonnell Foundation.

## Supporting Material

**Supporting Table 1A.** Average Event Rate (Hz) +/- standard error of the mean, and event count in parentheses, for the different physiological oscillation frequency bands. A1 Supra, A1 Gran, A1 Infra are from NHP A1 supragranular, granular, and infragranular sink channels, respectively. STG is from human iEEG recorded in supratemporal gyrus.

| Event Rate | Delta | Thet<br>a | Alph<br>a | Beta | Low<br>Gamma | Gam<br>ma | High<br>Gamma |
| --- | --- | --- | --- | --- | --- | --- | --- |
| **A1**<br>**Supra** | 0.36+<br>/-0.0<br>(10,600) | 0.56+<br>/-0.0<br>(16,544) | 0.66+<br>/-0.01<br>(19,476) | 1.54+<br>/-0.01<br>(44,920) | 1.29<br>+/-0.01<br>(37,800) | 4.44+<br>/-0.01<br>(130,088) | 13.12<br>+/-0.02<br>(383,848) |
| **A1**<br>**Gran** | 0.36+<br>/-0.0<br>(10,519) | 0.58+<br>/-0.0<br>(17,028) | 0.68+<br>/-0.0<br>(20,032) | 1.51+<br>/-0.01<br>(44,266) | 1.27<br>+/-0.01<br>(37,089) | 4.41+<br>/-0.01<br>(129,129) | 13.08<br>+/-0.02<br>(382,922) |
| **A1**<br>**Infra** | 0.36+<br>/-0.0<br>(10,563) | 0.57+<br>/-0.0<br>(16,756) | 0.64+<br>/-0.00<br>(18,874) | 1.43+<br>/-0.01<br>(41,912) | 1.23<br>+/- 0.01<br>(35,972) | 4.41+<br>/-0.01<br>(129,075) | 13.18<br>+/-0.02<br>(385,638) |
| **STG** | 0.37+<br>/-0.01<br>(820) | 0.55+<br>/-0.01<br>(1,204) | 0.74+<br>/-0.02<br>(1,630) | 1.50+<br>/-0.02<br>(3,287) | 1.28<br>+/- 0.02<br>(2819) | 3.29+<br>/-0.05<br>(7,246) | 12.26<br>+/-0.07<br>(26,917) |

**Supporting Table 1B.** Active time ratio (ATR) for the different physiological oscillation frequency bands. Mean value is presented (standard error was negligible). A1 Supra, A1 Gran, A1 Infra are from NHP A1 supragranular, granular, and infragranular sink channels, respectively. STG is from human iEEG recorded in supratemporal gyrus.

| ATR | Delta | Thet<br>a | Alph<br>a | Beta | Low<br>Gamma | Gam<br>ma | High<br>Gamma |
| --- | --- | --- | --- | --- | --- | --- | --- |
| **A1**<br>**Supra** | 0.44 | 0.29 | 0.19 | 0.25 | 0.13 | 0.26 | 0.34 |
| **A1**<br>**Gran** | 0.43 | 0.29 | 0.20 | 0.25 | 0.13 | 0.26 | 0.34 |
| **A1**<br>**Infra** | 0.44 | 0.30 | 0.19 | 0.23 | 0.13 | 0.25 | 0.34 |
| **STG** | 0.46 | 0.30 | 0.21 | 0.25 | 0.13 | 0.20 | 0.32 |

**Supporting Table 2-3** list ranges and mean+/-standard error of the mean.

**Supporting Table 2.** Cycles Per Event for the different physiological oscillation frequency bands. Range and mean+/-standard error of the mean, separated by semicolon (;). A1 Supra, A1 Gran, A1 Infra are from NHP A1 supragranular, granular, and infragranular sink channels, respectively. STG is human iEEG signals recorded from supratemporal gyrus.

| **Cycles Per Event** | **Delta** | **Theta** | **Alpha** | **Beta** | **Low Gamma** | **Gamma** | **High Gamma** |
| --- | --- | --- | --- | --- | --- | --- | --- |
| **A1 Supra** | 0.4-1<br>2.4;<br>2.8±-0.0 | 0.7-1<br>7.2;<br>3.3±-0.0 | 1.1-2<br>1.1;<br>3.5±-0.0 | 1.1-2<br>3.5;<br>3.6±-0.0 | 1.0-2<br>0.1;<br>3.6±-0.0 | 0.7-30.4;<br>3.7±-0.0 | 0.6-37.8<br>;<br>3.8±-0.0 |
| **A1 Gran** | 0.5-1<br>1.1;<br>2.8±-0.0 | 0.7-1<br>8.9;<br>3.3±-0.0 | 0.7-1<br>9.6;<br>3.5±-0.0 | 0.8-3<br>1.4;<br>3.7±-0.0 | 0.6-2<br>5.6;<br>3.66±-0.0 | 0.6-30.0;<br>3.7±-0.0 | 0.5-44.2<br>;<br>3.8±-0.0 |
| **A1 Infra** | 0.3-1<br>3.1;<br>2.8±-0.0 | 0.9-1<br>8.6;<br>3.4±-0.0 | 0.7-2<br>7.9;<br>3.5±-0.0 | 0.6-2<br>1.8;<br>3.6±-0.0 | 0.6-2<br>6.7;<br>3.6±-0.0 | 0.5-30.0;<br>3.6±-0.0 | 0.5-34.9<br>;<br>3.7±-0.0 |
| **STG** | 0.7-1<br>1.8;<br>3.0±-0.0 | 1.3-1<br>4.7;<br>3.7±-0.1 | 1.0-2<br>5.9;<br>3.5±-0.0 | 1.3-2<br>6.4;<br>3.8±-0.0 | 1.3-2<br>5.2;<br>3.73±-0.0 | 0.7-32.4;<br>3.24±-0.0 | 0.7-38.8<br>;<br>3.49±-0.0 |

**Supporting Table 3.** Number of local peaks in filtered waveforms for the different physiological oscillation frequency bands. Range and mean+/-standard error of the mean are presented, separated by semicolon (;). A1 Supra, A1 Gran, A1 Infra are from NHP A1 supragranular, granular, and infragranular sink channels, respectively. STG is human iEEG signals recorded from supratemporal gyrus.

| Local Peaks | Delta | Thet<br>a | Alph<br>a | Beta | Low<br>Gamma | Gam<br>ma | High<br>Gamma |
| --- | --- | --- | --- | --- | --- | --- | --- |
| **A1 Supra** | 0-13;<br>3.1+/-0.0 | 1-19<br>3.5+/-0.0 | 1-22;<br>3.6+/-0.0 | 0-28;<br>3.7+/-0.0 | 0-30;<br>3.7+/-0.0 | 0-42;<br>3.8+/-0.0 | 0-36;<br>3.8+/-0.0 |
| **A1 Gran** | 0-16;<br>3.2+/-0.0 | 1-26;<br>3.5+/-0.0 | 0-26;<br>3.6+/-0.0 | 0-30;<br>3.7+/-0.0 | 0-26;<br>3.7+/-0.0 | 0-32;<br>3.8+/-0.0 | 0-40;<br>3.8+/-0.0 |
| **A1 Infra** | 0-16;<br>3.2+/-0.0 | 0-26;<br>3.5+/-0.0 | 0-22;<br>3.6+/-0.0 | 0-31;<br>3.7+/-0.0 | 0-32;<br>3.7+/-0.0 | 0-46;<br>3.7+/-0.0 | 0-35;<br>3.8+/-0.0 |
| **STG** | 1-15;<br>3.2+/-0.1 | 1-21;<br>3.9+/-0.1 | 1-19;<br>3.6+/-0.0 | 1-25;<br>3.8+/-0.1 | 1-16;<br>3.5+/-0.0 | 1-26;<br>3.1+/-0.0 | 0-33;<br>3.4+/-0.0 |

**Supporting Table 4.** Logarithmic frequency bandwidth (Fspan) for the different physiological oscillation frequency bands. Values are mean+/-standard error of the mean. A1 Supra, A1 Gran, A1 Infra are from NHP A1 supragranular, granular, and infragranular sink channels, respectively. STG is human iEEG signals recorded from supratemporal gyrus.

| Fspan | Delta | Thet<br>a | Alph<br>a | Beta | Low<br>Gamma | Gam<br>ma | High<br>Gamma |
| --- | --- | --- | --- | --- | --- | --- | --- |
| **A1 Supra** | 0.580<br>+/-0.002 | 0.500<br>+/-0.002 | 0.490<br>+/-0.002 | 0.490<br>+/-0.001 | 0.480<br>+/-0.001 | 0.480<br>+/-0.001 | 0.470<br>+/-0.000 |
| **A1 Gran** | 0.580<br>+/-0.002 | 0.490<br>+0.002 | 0.480<br>+/-0.002 | 0.480<br>+/-0.001 | 0.480<br>+/-0.001 | 0.480<br>+/-0.001 | 0.470<br>+/-0.000 |
| **A1 Infra** | 0.590<br>+/-0.002 | 0.500<br>+/-0.002 | 0.490<br>+/-0.002 | 0.480<br>+/-0.001 | 0.480<br>+/-0.001 | 0.470<br>+/-0.001 | 0.470<br>+/-0.000 |
| **STG** | 0.600<br>+/-0.009 | 0.530<br>+/-0.007 | 0.500<br>+/-0.006 | 0.530<br>+/-0.004 | 0.490<br>+/-0.004 | 0.430<br>+/-0.003 | 0.470<br>+/-0.001 |

**Supporting Table 5.** Correlation between filtered and raw signals (Filter-match) for the different physiological oscillation frequency bands. Values are mean+/-standard error of the mean. A1 Supra, A1 Gran, A1 Infra are from NHP A1 supragranular, granular, and infragranular sink channels, respectively. STG is human iEEG signals recorded from supratemporal gyrus.

| Filter-match | Delta | Thet<br>a | Alph<br>a | Beta | Low<br>Gamma | Gam<br>ma | High<br>Gamma |
| --- | --- | --- | --- | --- | --- | --- | --- |
| **A1 Supra** | 0.400<br>+/-0.002 | 0.460<br>+/-0.001 | 0.470<br>+/-0.001 | 0.470<br>+/-0.001 | 0.490<br>+/-0.001 | 0.530<br>+/-0.000 | 0.620<br>+/-0.000 |
| **A1 Gran** | 0.400<br>+/-0.002 | 0.430<br>+/-0.001 | 0.450<br>+/-0.001 | 0.460<br>+/-0.001 | 0.490<br>+/-0.001 | 0.540<br>+/-0.000 | 0.630<br>+/-0.000 |
| **A1 Infra** | 0.410<br>+/-0.002 | 0.450<br>+/-0.001 | 0.450<br>+/-0.001 | 0.460<br>+/-0.001 | 0.480<br>+/-0.001 | 0.530<br>+/-0.000 | 0.630<br>+/-0.000 |
| **STG** | 0.200<br>+/-0.009 | 0.560<br>+/-0.008 | 0.580<br>+/-0.005 | 0.510<br>+/-0.003 | 0.390<br>+/-0.004 | 0.390<br>+/-0.003 | 0.260<br>+/-0.001 |

**Supporting Figure S6.**
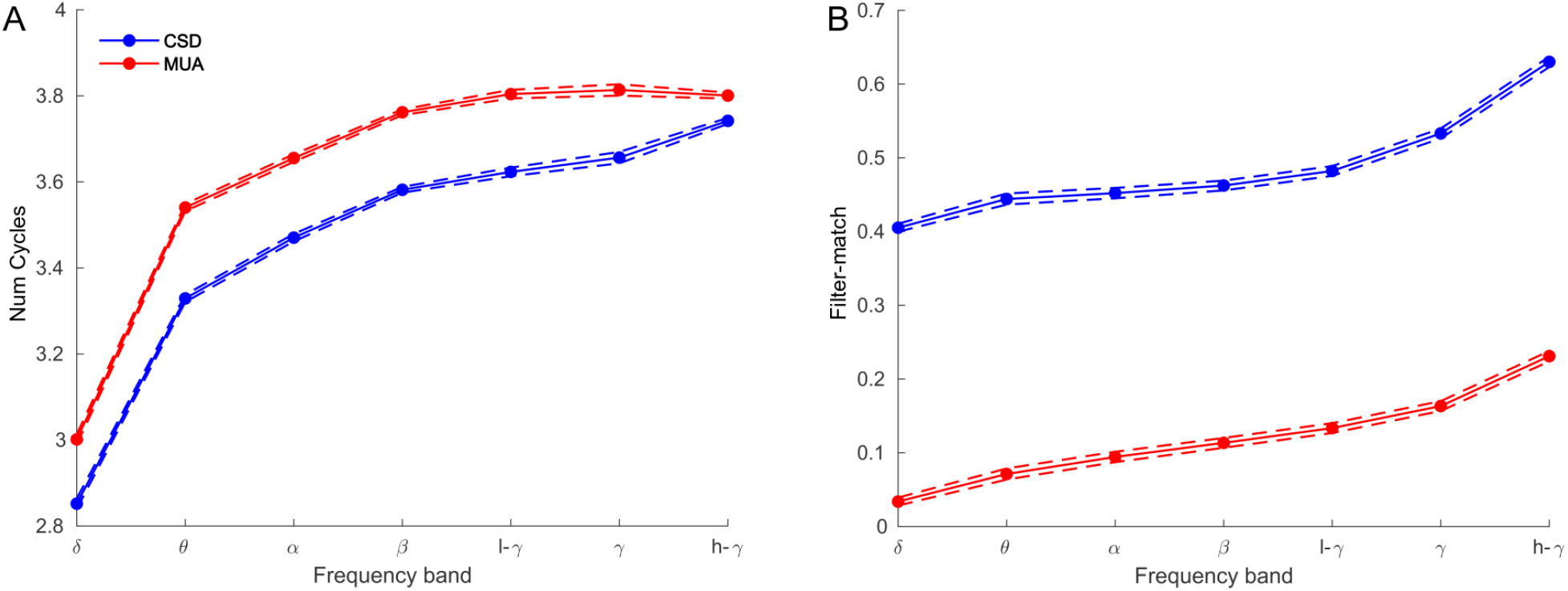
Comparison of features for oscillation events detected from NHP CSD (blue) and NHP multiunit activity (MUA; red). **(A)** While there is a higher number of cycles detected in MUA, the same pattern of increasing number of cycles with higher oscillation frequency is evident. **(B)** Filter-match was higher across the oscillation frequencies for CSD.

